# Tracking components of bilingual language control in speech production: an fMRI study using functional localizers

**DOI:** 10.1101/2023.02.07.527469

**Authors:** Agata Wolna, Jakub Szewczyk, Michele Diaz, Aleksandra Domagalik, Marcin Szwed, Zofia Wodniecka

**Author notes:** **Funding** This research was possible thanks to an OPUS grant from the National Science Center Poland, awarded to Zofia Wodniecka (2017/27/B/HS6/00959). **Declaration of Competing Interests** Authors declare no competing interests.

## Abstract

When bilingual speakers switch back to speaking in their native language (L1) after having used their second language (L2), they often experience difficulty in retrieving words in their L1: this phenomenon is referred to as the *L2 after-effect.* We used the L2 after-effect as a lens to explore the neural bases of bilingual language control mechanisms. Our goal was twofold: first, to explore whether bilingual language control draws on domain-general or language-specific mechanisms; second, to investigate the precise mechanism(s) that drive the L2 after-effect. We used a precision fMRI approach based on *functional localizers* to measure the extent to which the brain activity that reflects the L2 after-effect overlaps with the language network (Fedorenko et al., 2010) and the domain-general Multiple Demand network (Duncan et al., 2010), as well as three task-specific networks that tap into interference resolution, lexical retrieval, and articulation. Forty-two Polish-English bilinguals participated in the study. Our results show that the L2 after-effect reflects increased engagement of domain-general but not language-specific resources. Furthermore, contrary to previously proposed interpretations, we did not find evidence that the effect reflects increased difficulty related to lexical access, articulation, and the resolution of lexical interference. We propose that difficulty of speech production in the picture naming paradigm – manifested as the L2 after-effect – reflects interference at a non-linguistic level of task schemas or a general increase of cognitive control engagement during speech production in L1 after L2.

## Introduction

In their daily lives, bilinguals must face a constant challenge related to the non-selectivity or constant co-activation of their two languages (Costa et al., 1999; Kroll et al., 2006). To cope with this problem, bilinguals need an efficient system of control that resolves interference between the competing languages (Green, 1998), or a selection mechanism (Blanco-Elorrieta & Caramazza, 2021) that allows them to efficiently use the intended language. In this study we aimed to characterize the neurocognitive mechanisms of control that are engaged in bilingual speech production by capitalizing on the advantages of the functional localizer approach. To this aim, we explored how the control mechanisms affect bilingual speech production by looking at the *L2 after-effect*: a difficulty in native language (L1) production observed after prior use of a second language (L2).

### Bilingual language control: domain-general or domain-specific?

One of the most fundamental questions regarding the complex system that bilinguals use to control their two languages is whether it is based on a set of mechanisms specifically tailored to handle language (so called domain-specific, or language-specific mechanisms), or whether bilingual language control is achieved by domain-general cognitive control mechanisms (Abutalebi & Green, 2013; Wu et al., 2019; for a review see: Calabria et al., 2018).

So far, most evidence for the engagement of domain-general control mechanisms in bilingual language production comes from behavioral studies in which bilinguals were required to switch between their languages. The overall conclusion from these experiments is that switching between two languages incurs additional processing costs (Costa & Santesteban, 2004; Gollan & Ferreira, 2009; Meuter & Allport, 1999; Verhoef et al., 2009; for review see: Bobb & Wodniecka, 2013; Declerck et al., 2020; Declerck & Koch, 2022; Declerck & Philipp, 2015). This additional cognitive effort has also been observed as increased neural activation in structures associated with broadly defined cognitive control mechanisms (e.g., Abutalebi & Green, 2007; Coderre et al., 2016; De Baene et al., 2015; Wu et al., 2019; for reviews see: Abutalebi & Green, 2016; Luk et al., 2012; Sulpizio et al., 2020). It has been argued that bilingual language control is supported by domain-general mechanisms because task switching (which recruits domain-general control) and language switching share a number of computational processes, e.g., context monitoring, reconfiguring task-sets, engaging and disengaging from a task, suppressing the interference between competing representations and selective response inhibition (Green, 1998; Green & Abutalebi, 2013; Wu et al., 2019). This is also supported by research showing that the brain networks that support language switching and task switching overlap (Blanco-Elorrieta & Pylkkänen, 2016; Coderre et al., 2016; De Baene et al., 2015; Luk et al., 2012; Mendoza et al., 2021; Nee et al., 2007; Niendam et al., 2012; Weissberger et al., 2015; Wu et al., 2019).

On the other hand, the overlap between brain networks activated by language– and task-switching may not provide conclusive evidence for the domain-general nature of bilingual language control. First, language switching also engages brain structures that are not implicated in non-linguistic switching tasks (De Baene et al., 2015; Mendoza et al., 2021; Weissberger et al., 2015). Second, the overlap between the brain networks recruited by language– and task-switching may result from idiosyncratic properties of the switching task itself rather than bilingual language control. More specifically, it can be argued that language switching is an “artificial” task in the sense that it requires changing languages very frequently and depending on external cues, while switching in natural contexts is largely dictated by the speaker (i.e., internal cues; for a discussion on how cues affect switch costs, see Blanco-Elorrieta & Pylkkänen, 2017 for natural vs. artificial cues and Gollan et al., 2014 and Gollan & Ferreira, 2009 for voluntary vs. forced cues). As such, a language switching task may require the use of a very specific task set and keeping both languages co-activated. Such extra demands may recruit mechanisms engaged in performing tasks that are untrained, difficult, or not automated (Chein & Schneider, 2005; Crittenden & Duncan, 2014; Shashidhara et al., 2019), and these types of tasks are known to reliably activate the domain-general Multiple Demand network. What is more, bilinguals rarely switch between their languages on a word-by-word basis (Wodniecka et al., 2020; see also: Bobb & Wodniecka, 2013); and forced switching is related to greater engagement of cognitive control than voluntary switching (*for behavioral evidence see:* Gollan et al., 2014; Gollan & Ferreira, 2009; *for fMRI evidence see:* Reverberi et al., 2018; Zhang et al., 2015).

### Digging deeper – specific processes involved in bilingual language control

Besides the more general question of whether bilinguals rely on domain-general or language-specific networks when using their two languages, another unresolved issue is what specific processes are involved in bilingual language control. Models of bilingual language production assume that each language of a bilingual speaker is subserved by a set of different lexical, phonological, and articulatory representations (e.g., Blanco-Elorrieta & Caramazza, 2021; Costa, 2005; La Heij, 2005). Successful production requires using representations that belong to the target language but not to the non-target language. In the case of bilingual speakers (especially L1-dominant speakers), it is typically assumed that using their native language (L1) does not require additional control over representations of their second language (L2) because L1 representations are readily available and receive no interference from L2 representations. However, when an L1-dominant bilingual wants to use their weaker L2, the representations of L2 are less active and receive interference from strongly activated L1 representations, so accessing them may require additional control processes. The most prominent theory of bilingual language control proposes that resolution of between-language interference is implemented by active inhibition of L1 representations (Green, 1998); however, it does not specify at which level of word processing (e.g., lexical, phonological, articulatory) inhibition is implemented. On the other hand, other theoretical models argue that inhibition is not necessary for the successful selection of the target language (Blanco-Elorrieta & Caramazza, 2021; Costa, 2005; Koch et al., 2010). For example, the persisting activation account assumes that when a bilingual wants to use the weaker L2, its representations receive additional activation, which allows them to surpass the level of activation of the stronger L1 (for a review see: Koch et al., 2010). A recent extension of the activation-based account proposed that activation of target representations does not operate on a whole-language level (i.e., it does not apply to all items of L1 or L2) but it is rather a sum of activations driven by different factors, such as baseline word frequency, recency of use and communicative context (Blanco-Elorrieta & Caramazza, 2021). As such, both bilingual and monolingual speech production can be explained by the same selection mechanisms that select the most active representation at a given moment.

### The L2 after-effect as a window to study neurocognitive mechanisms of bilingual language control

As has been already indicated, the lion’s share of evidence on the involvement of control in bilingual language production comes from the language-switching paradigm (for a recent review see Sulpizio et al., 2020) which – as we argued above – has several important limitations. Some of these limitations are circumvented in a blocked language picture naming task that does not require as frequent switches between languages as the language switching task and yet allows control mechanisms involved in bilinguals’ language production to be tested. Several studies have observed that completing a task in L2 results in a subsequent slow-down of naming or an increase of errors committed in L1 (Branzi et al., 2014; Degani et al., 2019; Gollan & Ferreira, 2009; Van Assche et al., 2013; Wodniecka, Szewczyk, et al., 2020). This difficulty in using L1 after a short exposure to L2 has been referred to as the L2 after-effect. Any task involving language production in L2 can lead to the L2 after-effect: simply reading a list of words aloud (Degani et al., 2019), naming a set of pictures (Branzi et al., 2014, 2016; Gollan & Ferreira, 2009; Guo et al., 2011; Wodniecka et al., 2020) or coming up with words belonging to a given semantic category (Van Assche et al., 2013).

Available theories of bilingual speech production provide different explanations of the L2 after-effect. According to inhibition-based accounts, the retrieval difficulty that speakers experience in L1 after using L2 is a lingering consequence of the strong inhibition that had to be applied to L1 representations during L2 use (Green, 1998; Guo et al., 2011; Misra et al., 2012). Under the inhibitory accounts, resolution of the interference that drives the L2 after-effect relies on a set of domain-general mechanisms (e.g., Abutalebi & Green, 2016; Green, 1998; Green & Abutalebi, 2013). An alternative proposal assumes that the L2 after-effect is driven by increased interference between the representations of L1 and L2 due to the carry-over effects of increased L2 activation (Blanco-Elorrieta & Caramazza, 2021; Branzi et al., 2014, 2016; Koch et al., 2010).

To date, the origin of the L2 after-effect has been explored in three neuroimaging studies using fMRI. They revealed that the difficulty in word retrieval in L1 after using L2 results in increased engagement of brain regions often linked to domain-general control mechanisms (Branzi et al., 2016; Guo et al., 2011; Rossi et al., 2018) such as interference suppression (left middle frontal gyrus, left inferior parietal gyrus; Guo et al., 2010), visual attention shifts (left parietal cortex; Guo et al., 2010), articulation (right postcentral gyrus; Guo et al., 2010), response inhibition (right inferior frontal cortex; Branzi et al., 2016 right frontal pole and right superior parietal lobule; Rossi et al., 2018), response selection (left prefrontal cortex; Branzi et al., 2016), attention (bilateral inferior parietal lobules; Branzi et al., 2016), and conflict monitoring and resolution (bilateral anterior cingulate cortex; Rossi et al., 2018). While most of these structures have been linked to several different domain-general control mechanisms, some structures are thought to support language-specific processing as well as articulation. As such, the results of previous studies do not unanimously identify the mechanisms that drive the L2 after-effect.

### Capturing the neural basis of the L2 after-effect: methodological challenges

Importantly, three studies that explored the L2 after-effect were not free of methodological challenges. Two out of three studies report neural contrasts that involved different samples of participants (Guo et al., 2011; Rossi et al., 2018). This could be problematic as the engagement of language control mechanisms may be dependent on some aspects of language experience (see Casado et al., 2022 for evidence of the influence of language balance on the magnitude of engagement of global language control). What is more, a study by Rossi et al. (2018) compared a sample of bilinguals to monolinguals, but this does not make it possible to disentangle the effects of previous use of L2 from more general differences between bilingual and monolingual language processing. Branzi and colleagues (2016) addressed these limitations by comparing preceding context effects within the same bilingual participants; however, in this study, the baseline L1 block, which was compared to L1 after L2 in order to estimate the L2 after-effect, was in some participants preceded by a task in L2. Since the L2 after-effect is relatively long-lasting (at least 5 minutes, see Branzi et al., 2014; Casado et al., 2021; Wodniecka, Szewczyk, et al., 2020; for review see: Wodniecka, et al., 2020), a baseline for comparison (i.e., L1 naming) should not be preceded by a task in L2. It is therefore possible that the observed L2 after-effect was confounded by factors such as training, fatigue, or accumulating semantic interference (Runnqvist et al., 2012; for a discussion of how trial-related factors can affect the measurement of L2 after-effect see: Casado et al., 2022).

Moreover, the previous studies relied on group-level fMRI analysis. This approach focuses on the commonalities in activations between participants but is less sensitive to individual variability in brain organization, which is especially important among bilinguals whose variable language experience (related to a number of factors, such as immersion, age of acquisition, proficiency, daily use of language), may translate into changes in the functional and structural organization of the brain (e.g., DeLuca et al., 2019, 2020; Fedeli et al., 2021). What is more, while previous studies undoubtedly identified brain structures engaged in the L2 after-effect, the interpretations regarding the precise mechanisms represented by the observed brain response were not directly tested and were proposed based on the function assigned to a given anatomical structure (the so-called *reverse inference* problem, Poldrack, 2006; for a discussion of issues related to this approach in studying language see Fedorenko, 2021). It has been shown that macroanatomical cortical areas are characterized by vast inter-individual variability, which means that the link between anatomical structure and functional organization is not straightforward, especially for high-level cognitive processes such as language (Frost & Goebel, 2012; Tahmasebi et al., 2012; for a discussion see Fedorenko, 2021; Michon et al., 2022).

### Current study

In the current study, we explored the nature of bilingual language control by looking at the brain basis of the L2 after-effect. To address the limitations of experimental designs previously used to measure the L2 after-effect, we used a within-participant design that carefully controlled for the effects of the preceding language context as well as for possible effects of task order and fatigue. We had two specific goals: first, to determine to what extent the control mechanisms engaged in bilingual language control are domain-general vs. language-specific; second, to identify the specific process that may underlie difficulties in L1 production following L2 use, namely lexical retrieval difficulty (Casado et al., 2022), articulatory difficulty (Guo et al., 2011), and increased interference between languages (Branzi et al., 2014). We addressed these questions by using fMRI and a set of functional localizers, which – to the best of our knowledge – is an approach that has not been used before in studies exploring the mechanisms of language control in bilinguals. A crucial advantage of functional localizers is that they allow the modeling of brain responses at a subject-specific level; this increases the sensitivity of the statistical analyses and models inter-individual variability better than typical group level analyses.

To explore the domain-specificity or domain-generality of bilingual language control mechanisms engaged in the L2 after-effect, we used two well-established functional localizers which make it possible to track down subject-specific networks that correspond to the language system (Malik-Moraleda, Ayyash et al., 2022; Fedorenko et al., 2010) and the Multiple Demand network (Duncan, 2010; Fedorenko et al., 2013). The language localizer is a well-established paradigm that allows the identification of a robust network in the brain which spans the frontal, temporal, and parietal regions, which support the processing of language (e.g., Malik-Moraleda, Ayyash et al., 2022; Fedorenko, 2021; Fedorenko et al., 2010, 2011). This network has a robust specificity for language: it is engaged during reading (e.g. Fedorenko et al., 2010), listening (e.g. Pallier et al., 2003; Scott et al., 2017) and production (Hu et al., 2022; Liljeström et al., 2015), but it shows little engagement in non-linguistic tasks (Benn et al., 2021; Chen et al., 2021; Fedorenko et al., 2011; Ivanova et al., 2021) or social reasoning (Shain et al., 2022). Importantly, even though the language network is usually localized using a comprehension-based task it is equally sensitive to language production (Hu et al., 2020).

The Multiple-Demand localizer, on the other hand, allows the identification of another robust network in the brain which responds to increased task demands across many different types of tasks (Assem, Blank, et al., 2020; Braver et al., 2003; Cole & Schneider, 2007; Dosenbach et al., 2006; Duncan, 2010; Fedorenko et al., 2011, 2013; Miller & Cohen, 2001). Activity within this network has been shown to decrease with the increasing automatization of task performance (Chein & Schneider, 2005) and increase as a function of task complexity (Crittenden & Duncan, 2014; Fedorenko et al., 2013; Shashidhara et al., 2019), time pressure and reward (Shashidhara et al., 2019). Importantly, the frontoparietal Multiple Demand network shows little or no domain-specificity (Assem, Glasser, et al., 2020): it is robustly activated in a range of different linguistic and non-linguistic cognitive tasks (Fedorenko et al., 2013; Dosenbach et al., 2006; Shashidhara et al., 2019), including low-level perceptual tasks (Cole & Schneider, 2007).

While both the Language and the Multiple Demand networks show vast cross-subject variability (Fedorenko et al., 2010; Smith et al., 2021), they show a clear functional and topographic dissociation on a single-subject level (e.g., Blank et al., 2014; Diachek et al., 2020). As such, if brain activity corresponding to the L2 after-effect is found within one of these two networks, it will provide straightforward evidence for the engagement of domain-general or language-specific mechanisms.

To investigate the specific mechanisms that drive the L2 after-effect, we used 3 additional localizers that focused on lexical access (a localizer based on the Verbal Fluency task), articulation (a localizer based on a simple articularion task), and lexical interference (a localizer based on the Stroop task). Each of them tapped into one of the previously described mechanisms that possibly underlies the L2 after-effect. The Stroop task (Stroop, 1935) was used to capture brain structures engaged in the resolution of interference and inhibition of non-target responses in speech production (Diamond, 2013). An overlap between these structures and those involved in the L2 after-effect would provide an argument that the L2 after-effect is driven by increased interference between the two languages.

The verbal fluency task was used to identify brain regions that are sensitive to the difficulty of lexical access. As previously discussed, difficulties in lexical access seem to be one of the most parsimonious explanations of the L2 after-effect (Wodniecka, Szewczyk, et al., 2020; Casado et al., 2022). The verbal fluency task is thought to engage cognitive mechanisms that support word retrieval (Birn et al., 2010; Friesen et al., 2021; Shao et al., 2014). Specifically, the verbal fluency task was shown to tap into the efficiency of the lexical search and selection processes (Shao et al., 2014) as well as executive function mechanisms engaged in word retrieval (Friesen et al., 2021; Martin et al., 1994). Therefore, if L1 retrieval difficulty after L2 is due to L1 lexical access being hampered, then the brain activity corresponding to the L2 after-effect should overlap with the network identified using a verbal fluency localizer. Finally, verbal fluency performance has been shown to depend either on individual differences between subjects that are related to vocabulary size and working memory capacity, or on the mere speed of information processing (Unsworth et al., 2011). Therefore, a subject-specific localizer based on a verbal fluency task allows us to better capture the inter-individual variability in lexical retrieval between subjects.

Finally, to test the extent to which the L2 after-effect is driven by the articulatory difficulty of word production in L1, we used a localizer that allows us to identify brain regions specifically linked to articulation (Basilakos et al., 2018). This localizer was based on a contrast between articulating syllables and performing a simple motor sequence (finger tapping). An fMRI experiment which examined the neural basis of the L2 after-effect suggested that speaking in L1 after being exposed to L2 entails the engagement of additional articulatory resources (Guo et al., 2011). Moreover, more-recent behavioral evidence (Broos et al., 2018) suggests that greater difficulty in speaking L2 compared to L1 only affects the late articulatory stage; if so, it is also plausible that exposure to L2 results in articulatory difficulties for subsequent L1 use.

## Method

### Participants

Forty-two Polish-English late bilinguals took part in the study. One participant was excluded due to excessive head motion (>2mm) in the main fMRI task, resulting in a final sample of forty-one participants (31 females, 10 males, mean age = 23.29, SD = 3.24, range: 19-32). All participants were Polish native speakers who learned English as their second language. To participate in the experiment, they had to declare sufficient proficiency in English (B2 or higher) and obtain at least 20/25 points on the General English Test by Cambridge Assessment (mean score = 21.32, SD = 1.94). In the behavioral session, the participants’ proficiency in English was assessed again using LexTale (mean score = 71.58%, SD = 9.91%; Lemhöfer & Broersma, 2012). In addition, participants completed a questionnaire in which they rated their proficiency in all known languages with regards to reading, writing, listening, speaking, and accent; participants also gave information on their daily use of all languages and the age at which they started to learn each of them. Detailed information on the participants’ proficiency, age of acquisition, and daily use of languages can be found in Table 1. All participants gave written consent for participation in the study. The study was approved by the Ethics Committee of the Institute of Psychology of the Jagiellonian University concerning experimental studies with human subjects.

**Table 1.**
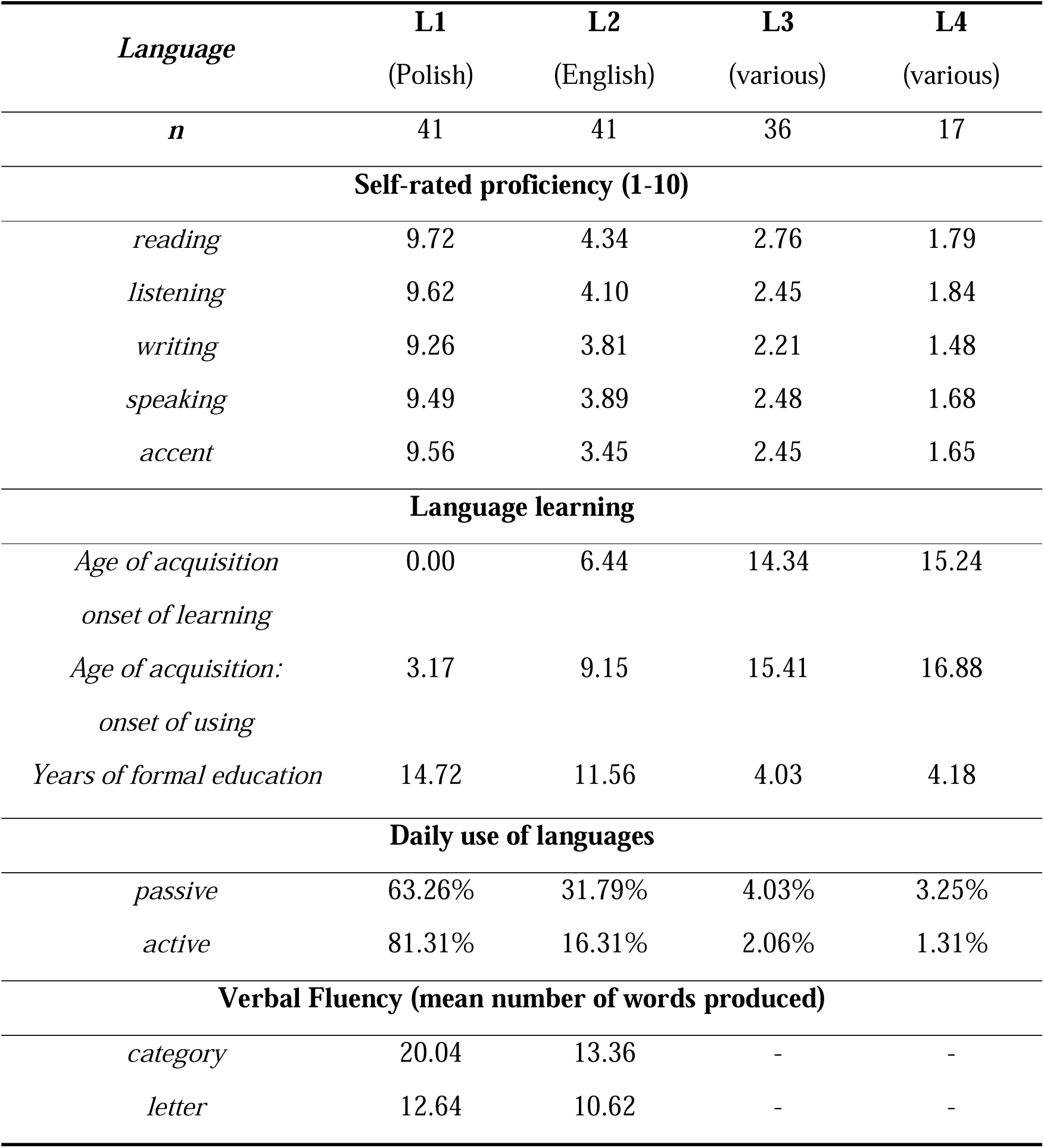
Language experience of participants. Information on self-rated proficiency, language learning and daily use of languages is given for all languages that participants declared to know. L1 always refers to Polish and L2 to English. L3 and L4 were a variety of different languages (incl. German, French, Italian, Spanish, Russian, Czech, Japanese, Korean, Norwegian, Latin and Esperanto). Not all the participants reported knowing an L3 or L4.

### Study Design

The experiment consisted of 3 testing sessions: one behavioral and two MRI scanning sessions. The design of the behavioral and MRI sessions is presented in detail in Figure 1, and the details of each task are discussed in the Design section.

**Figure 1.**
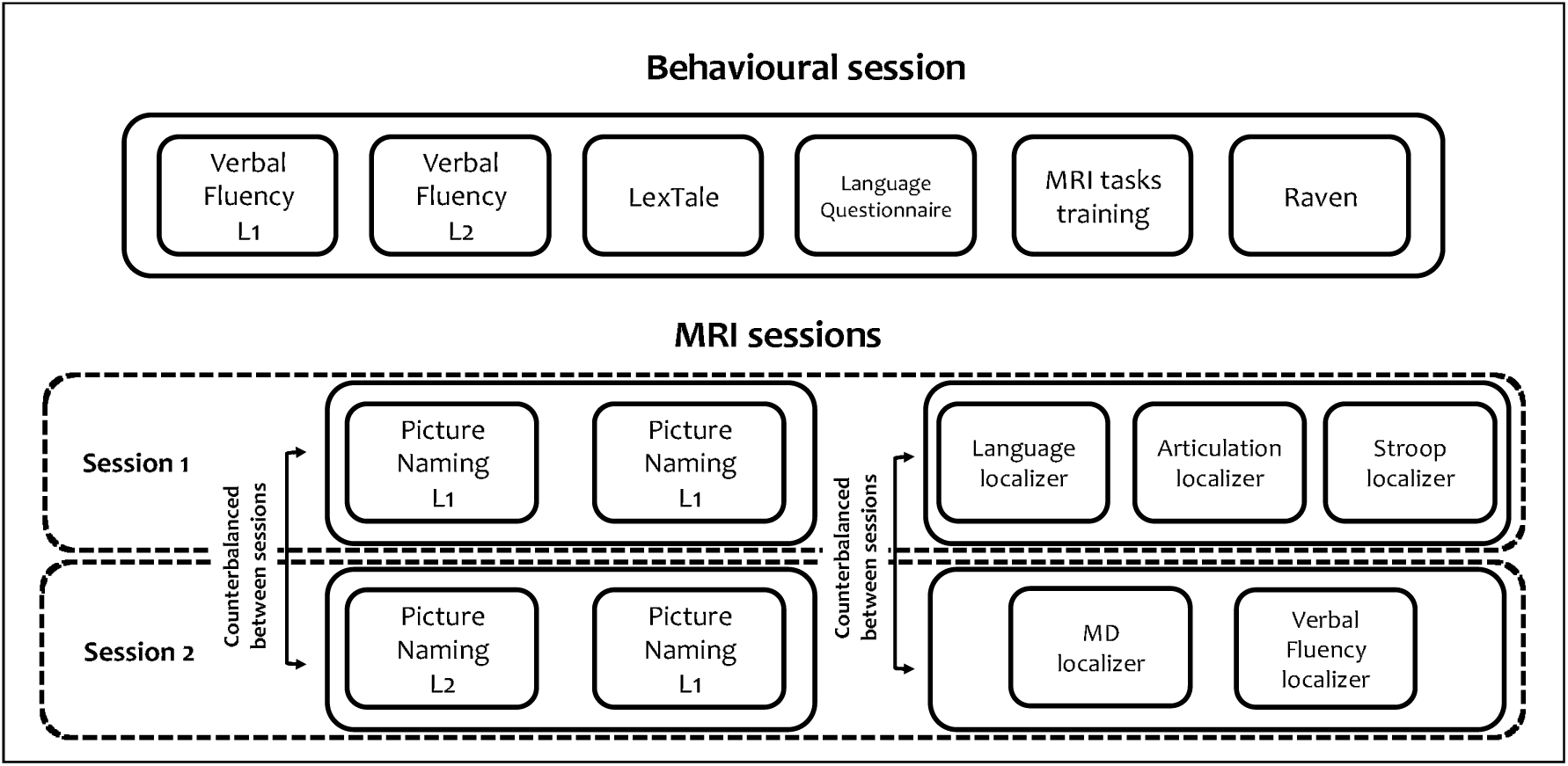
Summary of the design of the behavioral and MRI sessions. In the behavioral sessions, all participants completed the tasks in a fixed order. In the MRI sessions, the order of conditions completed in the first and the second session (experimental manipulation with L1 or L2) was independently counterbalanced between participants; the set of localizer tasks assigned to each session was also counterbalanced between participants.

In the behavioral session, participants completed a series of tasks, including semantic and phonetic verbal fluency in L1 and L2 (see details below), LexTale in English (Lemhöfer & Broersma, 2012), a language questionnaire, and a short version of Raven’s Progressive Matrices test (only uneven items) as a measure of a non-verbal intelligence. The behavioral session also included a short training session on the tasks to be performed in the MRI sessions.

During the MRI sessions, participants completed the main task, namely picture naming, and a series of localizer tasks. The main task was split between two sessions, with one condition (L1 or L2 in the first block) completed in the first session and the other condition completed in the second session, with the order counterbalanced across sessions. Localizer tasks included an auditory language localizer task (Malik-Moraleda, Ayyash et al., 2022; Scott et al., 2017), an articulation localizer, a Stroop task, a visual working memory task corresponding to the Multiple Demand localizer (Fedorenko et al., 2013), and a verbal fluency task (order counterbalanced, see Figure 1 for details). In tasks requiring overt spoken responses (i.e., picture naming, Stroop task, articulation task, and verbal fluency) participants’ responses were recorded with FOMRI III, which is an MR-compatible, fiber-optic microphone system (OptoacousticsLtd., Or-Yehuda, Israel). Participants were instructed to minimize jaw movements while giving the spoken responses, which they had practiced during the behavioral session.

### Materials and Procedure

#### Behavioral session

***Verbal Fluency task*.** In this task, participants were asked to produce as many words belonging to a given category (fruits or animals) or beginning with a given letter (B or M) as they could within a 1-minute time limit. Cues were counterbalanced between languages. Each participant first completed the task in Polish (with one category and one letter) and then in English (also with one category and one letter). Each cue (letter or category) was displayed on the screen for 1 minute, during which participants provided verbal responses that were recorded with the microphone. This task was used as a complementary measure of relative proficiency in L1 and L2; the mean number of words produced for letter and category tasks in each language is reported in Table 1.

***LexTale.*** In the LexTale task, participants were instructed to say whether the string of letters presented on a screen corresponded to an existing word in English. The response (yes/no) was provided via a keyboard. Each word or nonword appeared in the center of the screen and remained there until the participant responded. The task used the same words and nonwords as the original LexTale task (Lemhöfer & Broersma, 2012).

***Language questionnaire.*** In the language questionnaire, participants were asked to respond to four categories of questions. The first category concerned proficiency in the native and foreign languages that the participants knew. It included a self-rating scale on the five language skills (reading, writing, listening, speaking, and accent) and questions concerning the age of acquisition. The second category included questions on patterns of language use: the participants were asked to estimate how much time during a typical day they actively (i.e., speaking, writing) and passively (i.e., watching movies, reading, listening to music/radio/podcasts) use the languages they know. In the third category, participants were asked to provide information about language use contexts: home, work, school, and free time. Specifically, they were asked to estimate the number of hours that they use each language they know in each of these contexts. Finally, the fourth category of questions referred to demographic information, such as age, gender, and education. Filling out the entire questionnaire took approximately 15-20 minutes.

***MRI task training.*** In this part of the behavioral session, participants completed short versions of the tasks that they were going to perform during the MRI session (for details of the tasks see the MRI session design). Each task lasted approximately 1-2 minutes and was repeated if a participant asked for more practice. In tasks requiring overt responses, participants were asked to practice minimizing jaw movements while speaking.

***Raven test.*** In this task, participants completed a shortened version of Raven’s Advanced Progressive Matrices test (only odd-numbered items). They had 20 minutes to complete as many matrices as they could. The test was used in a paper-pencil format and was aimed to provide a measure of fluid intelligence.

#### MRI sessions – the main task

In the main task, participants were asked to overtly name pictures of objects in four task blocks: two blocks in each session. One task block corresponded to one functional run. The first block of each session corresponded to one of the experimental manipulations: exposure to L1 (which required naming pictures in L1) or exposure to L2 (which required naming pictures in L2). The order of experimental manipulation blocks was counterbalanced between participants, i.e., some of them started with the L2 exposure conditions in the first session and completed the L1 exposure condition in the second session; some of them started with the L1 exposure condition in the first session and completed the L2 exposure condition in the second session. In the second block of each session, participants were asked to name pictures in L1. Each block contained 55 colored pictures representing objects (from the CLT database: Haman et al., 2015; Wolna et al., 2022). Pictures were not repeated between blocks. Each block started with a fixation cross (“+”) displayed on a screen for 7000ms. Subsequently, 55 pictures were presented following a rapid event-related design. Each picture was displayed on a white background for 2000ms and was followed by a fixation cross, jittered at an interstimulus interval ranging from 607ms to 10144ms (M = 3091ms, SD = 2225ms). The order of picture presentation and interstimulus intervals was optimized using optseq2 (Dale, 1999). Finally, at the end of each block, a fixation cross was presented for 8000ms. Overall, each block of the Picture Naming task lasted 295s.

#### MRI sessions – localizer tasks

After completing the main task, in each session participants completed two or three localizer tasks. Each localizer consisted of two functional runs.

**Language localizer.** Participants listened to fragments of *Alice in Wonderland* in L1 and distorted speech recordings in which it was impossible to understand what the speaker was saying. The task followed the design introduced by Scott et al. (2017) and Malik-Moraleda, Ayyash et al. (2022), in which the localizer contrast was based on the intact > distorted speech condition contrast. The intact condition consisted of short passages from Alice in Wonderland read by the native speaker. The degraded speech condition was created based on the intact condition but, it sounded like poor, indecipherable radio reception of speech with preserved prosody but not decipherable words or phonemes. In each functional run, participants listened to 6 short passages of Alice in Wonderland in their native language (18s each) and 6 passages of distorted speech (18s each). Additionally, 4 fixation blocks (12s each) were included in each run. The total duration of each run was 264s. Each participant completed 2 runs of the task.

**Articulation localizer.** Participants performed two tasks: an articulation task in which they were asked to repeat out loud a string of previously memorized syllables (“bła” [bwa], “sza” [[a], “tra” [tra] – all frequently occurring in Polish as word constituents, but meaningless in isolation) and a motor task in which they were asked to perform a simple motor sequence with the fingers of both hands (touching the thumb with their index, ring, and little finger, respectively). The localizer was based on the articulation > motor sequence contrast. In each functional run, participants completed 6 articulation and 6 finger tapping blocks (18s each). Additionally, 4 fixation blocks (12s each) were included in each run. Each participant completed 2 runs of the task.

**Stroop localizer.** Participants were asked to overtly name the color of the presented word’s font. The design of the task followed the one used by Fedorenko et al. (2011), in which participants completed blocks of the easy and hard versions of the task. In the easy condition, the words corresponded to non-color adjectives [examples in the original language mased for review]; in the hard condition, the words corresponded to color adjectives [examples in the original language mased for review]. In the hard condition, half of the trials were conflict trials in which the word’s font color did not match its meaning; the other half were congruent trials. The order of trials within the hard condition was randomized. Each run started with a fixation cross (“+”) displayed on a gray background for 8000ms. Following that, participants completed 16 task blocks (8 per condition, 18s each). Task blocks contained 12 trials which started with a fixation cross displayed for 500ms followed by a word displayed for 1000ms. Additionally, five fixation blocks (18s) were included in each run. The order of experimental and fixation blocks was counterbalanced between participants and runs (four counterbalance lists were created and each participant completed two of them). The localizer was based on the hard > easy condition contrast. The total duration of one run was 394s. Each participant completed 2 runs of the task.

**Multiple Demand localizer.** Participants were asked to perform a spatial working memory task (e.g., Fedorenko et al., 2011, 2013). The design of this task followed the one used by Fedorenko et al. (2011), in which the localizer contrast was based on the hard > easy condition contrast. In each trial, participants saw four 3 x 4 grids and were asked to memorize the locations of the fields that were marked as black on each of the grids. In the easy condition, they had to memorize one location per grid (four in total); in the hard condition, they had to keep track of two locations per grid (eight in total). After memorizing the grids, participants had to choose a grid that contained all the memorized locations from among two grids presented on the screen. Each trial started with a fixation cross displayed on the screen for 500ms; this was followed by four grids, each displayed for 1000ms. After that, the choice task appeared on the screen until a response was given (for a max of 3750ms); this was followed by feedback (right or wrong) displayed for 250ms and a fixation cross, which was displayed for 3750ms minus the reaction time in the choice task. In each of the conditions (easy and hard) participants completed 4 trials per block (34s in total). Each run contained five easy condition blocks and five hard condition blocks. Additionally, six fixation blocks (16s) were included in each run. The order of experimental and fixation blocks was counterbalanced between participants and runs (four counterbalance lists were created and each participant completed two of them). The total duration of one run was 438s.

**Verbal fluency localizer.** Within 10 s blocks, participants were asked to produce as many words as they could that either pertained to a given category (semantic fluency) or started with a given letter (phonemic fluency). The entire task was completed in L1. In each run, participants completed six semantic fluency and six phonemic fluency blocks, each focusing on a separate letter or semantic category. Each letter and category were presented only once, and they did not repeat between blocks and runs. A full list of stimuli used in this task is available in the Supplementary Materials. In addition, each run included five baseline blocks, in which participants were asked to enumerate the months starting from January (Birn et al., 2010). The localizer contrast was based on task (semantic or phonemic fluency) > baseline condition contrast. Following Birn et al. (2010), each block was followed by a fixation cross displayed for 7s–13 s. Each run lasted 348s. Each participant completed 2 runs of the task.

### MRI Data Acquisition

MRI data were acquired using a 3T scanner (Magnetom Skyra, Siemens) with a 64-channel head coil. High-resolution, whole-brain anatomical images were acquired using a T1-weighted MPRAGE sequence (208 sagittal slices; voxel size 0.9×0.9×0.9 mm^3^; TR = 1800ms, TE = 2.32ms, flip angle = 8°). A gradient field map (magnitude and phase images) was acquired with a dual-echo gradient-echo sequence, matched spatially with fMRI scans (TE1 = 4.92ms, TE2 = 7.3ms, TR = 503ms, flip angle = 60°). Functional T2*-weighted images were acquired using a whole-brain echo-planar (EPI) pulse sequence (50 axial slices, 3×3×3 mm isotropic voxels; TR = 1400ms; TE = 27ms; flip angle = 70°; FOV 192 mm; MB acceleration factor 2; in-plane GRAPPA acceleration factor 2 and phase encoding A>>P) using 2D multiband echo-planar imaging (EPI) sequences from the Center for Magnetic Resonance Research (CMRR), University of Minnesota (Xu et al, 2013). In order to account for magnetic saturation effects, the first four volumes of each sequence were not included in the analysis.

### MRI Data preprocessing

All data were visually inspected for artifacts. The non-brain tissue was removed using the FSL Brain Extraction Tool (Smith, 2002). For the preprocessing of functional data, we used FSL FEAT (version 6.0.0; Woolrich et al., 2004). The preprocessing steps included high-pass temporal filtering (100s), spatial smoothing (FWHM = 5 mm), distortion correction of functional images using fieldmap with FUGUE, co-registration and normalization using FLIRT (Jenkinson et al., 2002; Jenkinson & Smith, 2001) and motion correction using MCFLIRT (Jenkinson et al., 2002) with 6 rigid-body transformations. Functional images were co-registered with their anatomical images; subsequently, they were registered to MNI standard space (FSL’s 2mm standard brain template).

### MRI Data analysis

In the 1^st^-level statistical analysis, a double-gamma hemodynamic response function was used to model the BOLD signal corresponding to each event (trial in the main task, as it followed the event-related design; and blocks in the localizer tasks which followed a blocked design). The estimates of motion obtained with MCFLIRT (Jenkinson et al., 2002) were included in the 1^st^-level GLM as nuisance covariates. For the localizer tasks, the two functional runs of each localizer task were combined in a second-level analysis for each participant using an FSL fixed-effect model. To ensure better comparability of our findings with previous studies, for language, Multiple Demand, and articulation localizers we used sets of functional parcels established and validated by previous studies instead of creating group-level parcels based on our data (twelve language parcels: Fedorenko et al., 2010; twenty multiple demand parcels: Fedorenko et al., 2013; eleven articulation parcels: Basilakos et al., 2018). It is important to note that the group-level parcels from previous studies showed a high overlap with parcels for the language, Multiple Demand, and articulation networks defined based on our data. For the remaining Stroop and Verbal Fluency localizers, group-constrained subject-specific ROIs were defined for each participant following the approach proposed by Fedorenko et al. (2010). In the first step, the individual subject activation maps thresholded at 5% of most active voxels. In most cases this threshold was slightly more conservative than z > 3.1, uncorrected at the whole brain level. We chose to use the 5% of most active voxels instead of a standard threshold because it yielded maps with comparable number of voxels between subjects that consequently produced more coherent group-level parcels. Subsequently, the individual parcels were binarized and overlaid on top of each other. The probabilistic map obtained this way was then thresholded so that only voxels active in more than 5 participants (∼20%) would be included. In the second step, the probabilistic maps were smoothed using an 8×8×8 Gaussian kernel and subsequently, group-level partitions were created using a watershed algorithm for segmentation. This procedure yielded nineteen group-level parcels for the Stroop task and fifteen group-level parcels for the Verbal Fluency task. Before further analyses, we excluded very small parcels (mean size across subjects smaller than 10 voxels). This yielded a final set of thirteen parcels within the Stroop network and thirteen parcels within the verbal fluency network. In each of these parcels significant activation to the localizer contrast, i.e.: color words > neutral words for the Stroop task and fluency > automated speech for verbal fluency (thresholded at z>2, corrected for multiple comparison using a cluster-wise correction at p<0.05) was found in at least 51% of participants for the Stroop parcels (mean = 70.36%) and in at least 73.17% of participants for the Verbal Fluency parcels (mean = 84.99%). In the last step, group-level partitions were intersected with subject-specific data, thus yielding a set of subject-specific group-constrained ROIs. Finally, we selected the top 10% most responsive voxels within each parcel based on the *z*-value map for a given localizer contrast (Language: speech > degraded speech, MD: hard > easy visual working memory task, Articulation: syllables > motor sequence, Stroop: color words > neutral words, and Verbal Fluency: fluency > automated speech) and we binarized the obtained ROIs. We then used the ROIs corresponding to all localizer tasks to extract parameter estimates corresponding to % signal change in the two critical conditions of the main task: naming in L1 after L1, and naming in L1 after L2. Parameter estimates were extracted from the 1st-level analyses for each participant separately. For the analysis of the functional localizer scope, we used an across-run cross-validation procedure to extract the % signal change corresponding to each localizer task condition (i.e., we defined the functional parcels based on the first functional run and estimated responses to the localizer contrast on the second run and repeated this procedure using the second run to define the functional parcels and the first run to extract response estimates; see Fedorenko et al., 2010). This was done using FSL’s FEATquery tool (http://www.FMRIb.ox.ac.uk/fsl/feat5/featquery.html). Further analyses were performed on the extracted parameter estimates in R (version: 4.0.2; R Core Team, 2020). Following an approach that proposes modeling activity within a functional network on the network level instead of relying on separate analyses for each ROI (Malik-Moraleda et al., 2021), we fitted one linear mixed-effect model for each functional network according to the following formula:

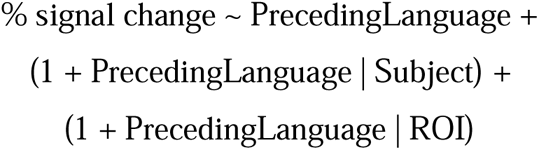

In each model, the fixed effect of PrecedingLanguage corresponded to a predictor with two levels: *L1 after L1* and *L1 after L2*. Before the analysis, this categorical predictor was deviation-coded (L1 after L1 = –0.5; L1 after L2 = 0.5). The analysis was performed using the lmer() function from the lmerTest package (Kuznetsova et al., 2017)For each functional network, we first fitted a maximal model and then identified the best random-effects structure following the recommendations of Bates et al. (2018). The reason for modeling the activity within each network with ROI as a random effect instead of a fixed effect relies on the assumption that the different brain regions engaged within a given functional network exhibit a significant degree of functional integration. To further explore the fine-grained differences between our critical conditions, which might only affect a subset of functional parcels within each network, we also fitted separate models for each ROI in each of the functional networks. The results of the ROI-specific analyses are reported in the *Supplementary Materials*. Raw neuroimaging data used in this study are freely available at the OpenNeuro repository https://openneuro.org/datasets/ds004456. Data and code necessary to reproduce the ROI and behavioral analyses are freely available at https://osf.io/59za8/?view_only=e23f3500d12e4109b75ca18fdbd320e5.

## Results

### Functional localizers’ scope

Functional networks identified with a set of localizer tasks were based on robust contrasts that yielded strong effects in each network (language localizer: intact vs. degraded speech: β = 0.520, *t(40)* = 5.684, *p* < .001; Multiple Demand localizer: hard vs. easy visual working memory task: β = 0.580, *t(40)* = 13.12, *p* < .001; Articulation localizer: articulation vs. motor task: β = 1.337, *t(40)* = 20.84, *p* < .001; Stroop localizer: Stroop task blocks vs. congruent color naming: β = 0.435, *t(40)* = 11.75; *p* < .001; Verbal Fluency localizer: fluency vs. enumerating months: β = 1.155, *t(40)* = 16.73, *p* < .001). The localizer tasks yielded five sets of parcels covering a wide range of brain regions. Language and Multiple Demand parcels covered two robust non-overlapping networks in the brain (see Figure 2A, although small parts of language and Multiple Demand network show some overlap, this was only the case on for the group-level parcels, not the individual subject-specific parcels). The Articulation parcels included a set of regions partially overlapping with the language network in the temporal ROIs but it also included parcels that selectively responded to articulation but not language (see figure 2B). As expected, the Stroop network largely overlapped with the Multiple Demand network (for similar results see Fedorenko et al., 2013), however, the Stroop parcels were considerably smaller than the Multiple Demand network (see figure 2C). The differences in size between the Stroop and the Multiple Demand networks are partially driven by the fact that the Stroop parcels were created based on single-subject activation maps thresholded at 5% of the most active voxels (for details see Methods), which puts an a priori constraint on the size of the activation maps and the parcels that are created based on these maps. However, the Stroop network was also considerably smaller than the Multiple Demand network when other, less stringent threshold were used (e.g., z>2). As such, the Stroop network can be considered a functional sub-system of the larger and more general Multiple Demand network. Finally, the Verbal Fluency localizer yielded a set of strongly left-lateralized parcels partially overlapping both with the Language and the Multiple Demand networks (for Verbal Fluency parcels see Figure 4).

**Figure 2.**
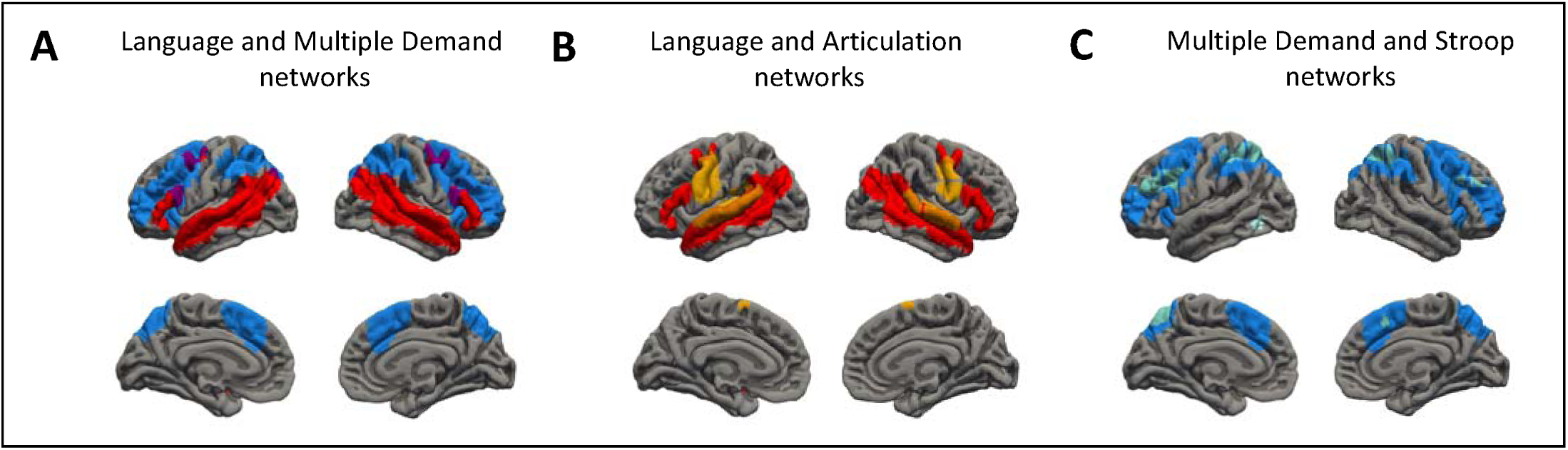
Brain masks corresponding to group-level functional networks. The figure presents a comparison of brain networks identified using different functional localizers (A) Language (red) and Multiple Demand (blue) networks; (B) Language (red) and articulation (yellow) networks; (C) Multiple Demand (blue) and Stroop (turquoise) networks. All networks presented on the graph correspond to a combination of group-level parcels.

To further characterize the functional networks as well as the scope of the localizer tasks used in this study, we ran additional analyses showing (1) responses to all localizer tasks used in this study within the language and the Multiple Demand networks; and (2) responses to the Language and Multiple Demand localizer tasks (i.e., listening to degraded vs. intact speech and hard vs. easy working memory task) in all five functional networks used in this study. Results of these analyses can be found in the Supplementary Materials.

### Bilingual language control: domain-general or language-specific

To address the question of whether the bilingual language control mechanisms driving the L2 after-effect rely on domain-general or domain-specific mechanisms, we compared brain activity corresponding to naming pictures in *L1 after L1* and *L1 after L2* within two functional networks: the language network and the Multiple Demand network. The results revealed no significant effect of the preceding language within the language network (β = 0.026, *t(40)*= 0.89, *p* = .379, effect size = 0.28); however, a significant effect of the preceding language was found within the Multiple Demand network (β = 0.072, *t(40)* = 2.679, p = .011, effect size = 0.85). The results of the comparison between *L1 after L1* and *L1 after L2* within the language and Multiple Demand networks are presented in **Figure 3**.

**Figure 3.**
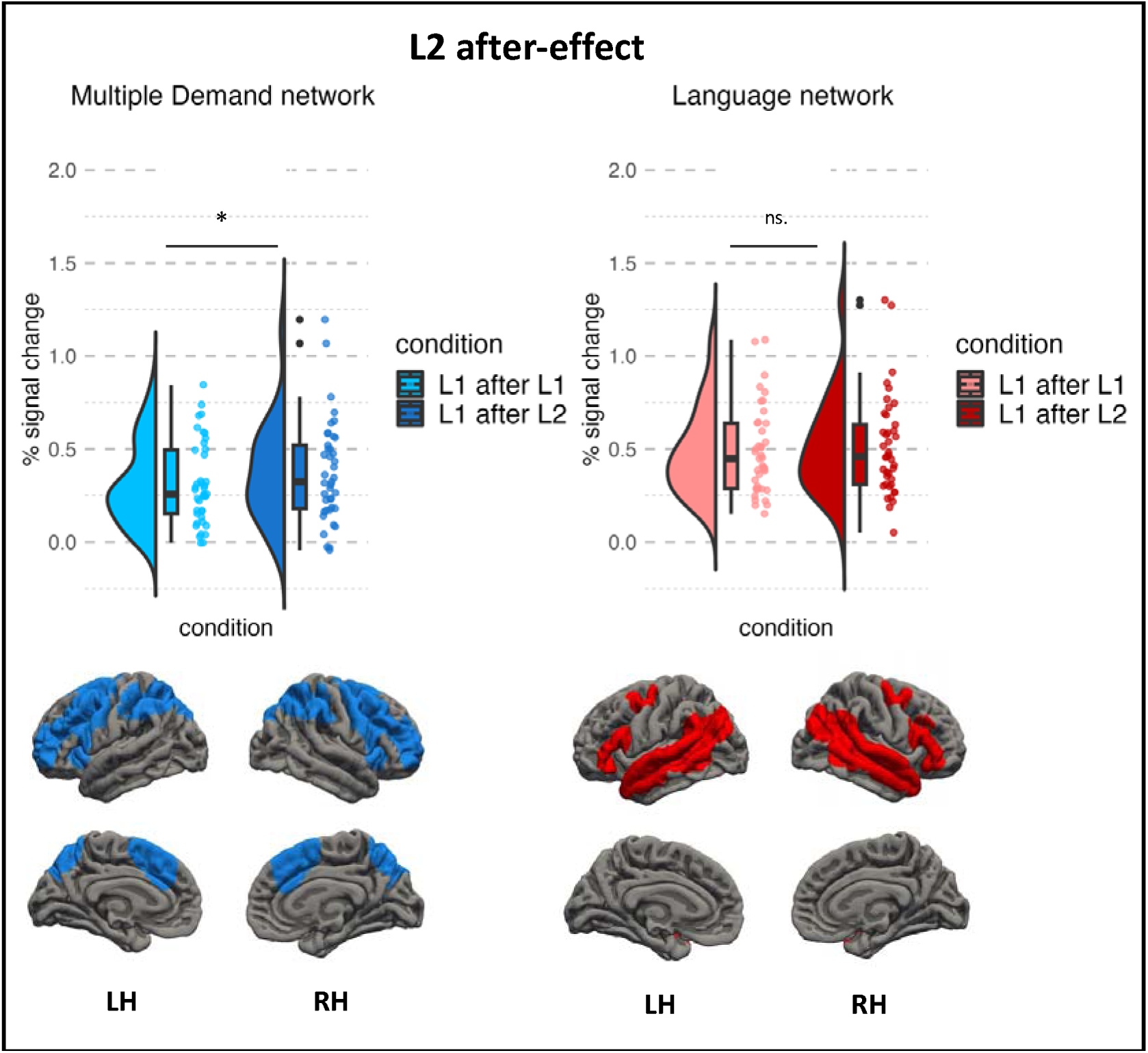
Neural activation corresponding to the L2 after-effect within the Multiple Demand and Language networks. Plots represent % BOLD signal change in response to naming pictures in L1 after L1 (light hues) and L1 after L2 (dark hues). Rendered maps present sets of group-level parcels used in the subject-specific analysis (LH = left hemisphere; RH = right hemisphere).

Separate analyses on each individual ROI showed significant differences between *L1 after L1* and *L1 after L2* in four ROIs in the Multiple Demand network (left and right angular gyrus / superior parietal lobule, right superior and middle frontal gyrus, and right frontal pole), but no significant results were found in the language network. Full results showing comparisons between conditions for each ROI and each network are presented in **Tables S1 and S2** in the *Supplementary Materials*.

### Digging deeper: specific mechanisms of bilingual language control

To explore which specific mechanisms drive the L2 after-effect, we compared the brain activity corresponding to naming pictures in *L1 after L1* and *L1 after L2* within three functional networks identified with the Articulation, Stroop, and Verbal Fluency tasks. We did not find a significant effect of the preceding language in any of the three networks (Articulation: β = 0.005, *t(40)* = 0.09, *p* = .926, effect size = 0.03; Stroop: β = 0.034, *t(40)* = 1.02, *p* = .314, effect size = 0.32; Verbal Fluency: β = 0.003, *t(40)* = 0.07, *p* = .941, effect size = 0.02; see Figure 4). Separate analyses on each individual ROI for each of these networks are presented in **Tables S3-5** in the *Supplementary Materials*.

**Figure 4.**
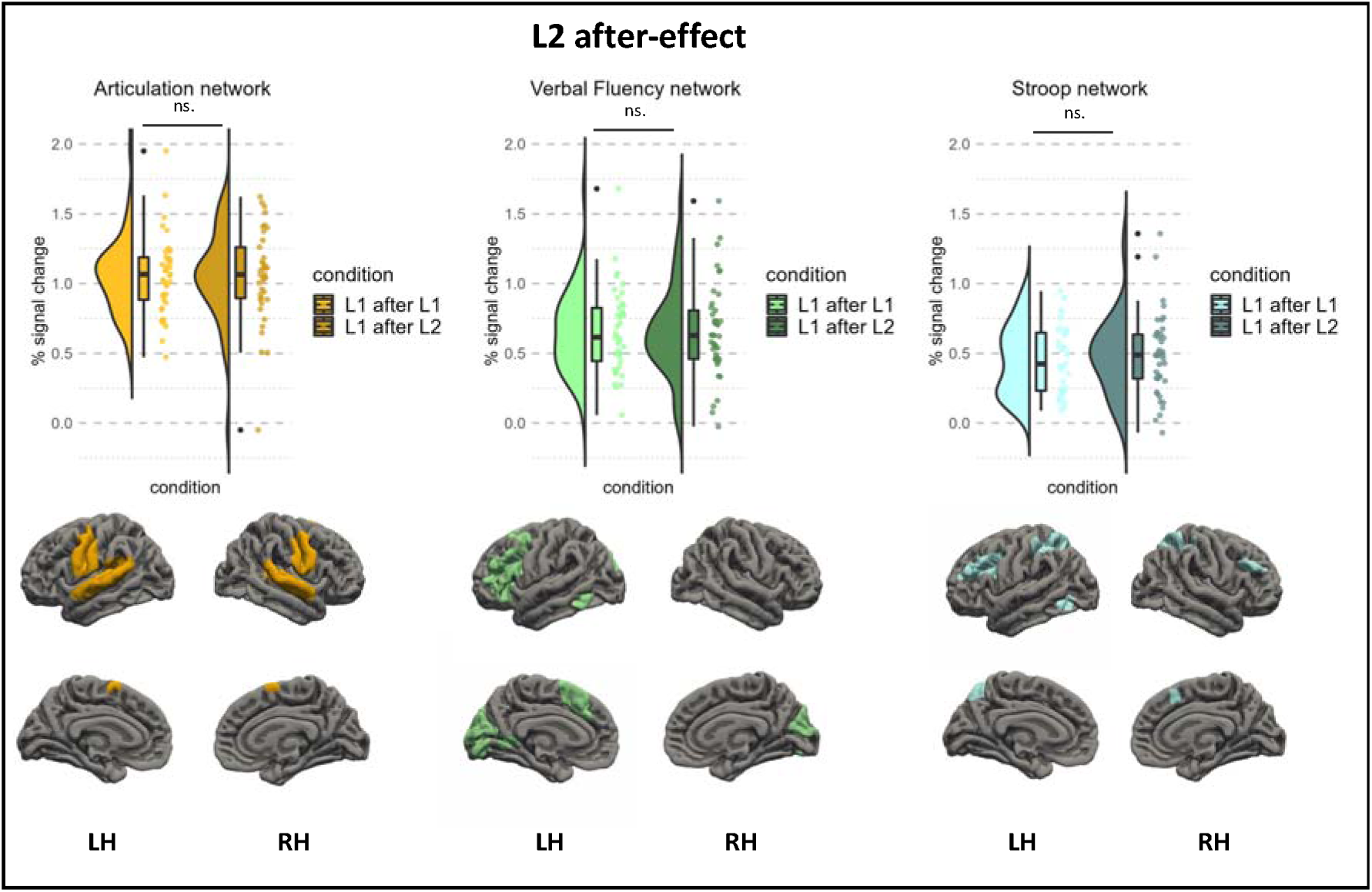
Neural activation corresponding to the L2 after-effect within the Articulation, Verbal Fluency and Stroop networks. Plots represent % BOLD signal change in response to naming pictures in L1 after L1 (bright hues) and L1 after L2 (dark hues). Rendered maps present sets of group-level parcels used in the subject-specific analysis (LH = left hemisphere; RH = right hemisphere).

## Discussion

### Bilingual language control: domain-general or language-specific?

We explored the neurocognitive mechanisms underlying the L2 after-effect: a difficulty in producing words in the native language (L1) after having used the second language (L2). We compared brain activity corresponding to speaking in L1 after L2 with speaking in L1 after L1 within functionally defined brain networks. Our goal was to test (1) whether the L2 after-effect is related to activity in domain-general or domain-specific brain networks, and (2) whether the L2 after-effect reflects lexical retrieval difficulty, interference resolution, or increased difficulty at the phonological and articulatory stages of speech production. Unlike previous work exploring the neural basis of bilingual speech production, we used individual-level functional localizers (Fedorenko et al., 2010) to provide functional interpretations of brain responses associated with the L2 after-effect.

To address the question of whether the L2 after-effect is associated with increased engagement of language-specific and/or domain-general brain networks, we compared brain responses to speaking in L1 after L2 and L1 after L1 with two well-established, robust functional networks: the language network (Malik-Moraleda, Ayyash et al., 2022; Fedorenko et al., 2010; Scott et al., 2017) and the Multiple Demand network (Duncan, 2010; Fedorenko et al., 2013). We found that the difficulty of naming pictures in L1 after L2 is related to increased BOLD responses within the domain-general Multiple Demand network, but not within the domain-specific language network. Furthermore, to explore whether the L2 after-effect is driven by lexical access difficulty, increased interference between language-specific representations, or articulatory difficulty, we used three additional functional tasks as localizers: verbal fluency, the Stroop task, and an articulation task. We found that the L2 after-effect does not overlap with any of these process-specific localizers. We have also replicated the results of a behavioral study (Casado et al., 2022) showing that the magnitude of the L2 after-effect depends on the balance between languages, defined as a difference between mean naming latencies in L2 and L1 (see *Supplementary Materials* for a detailed report). This is an important note as our study is the first to record naming latencies corresponding to the L2 after-effect in the scanner. As such, our data provides straightforward evidence that the L2 after-effect can be successfully tested in the MRI scanner, despite the noisy and challenging environment.

### The L2 after-effect reflects the increased engagement of domain-general cognitive control

We found evidence for increased response for speech production in L1 after L2 compared to L1 after L1 in the Multiple Demand network but not within the language network (the interaction between condition and network was, however, not significant (β = 0.046, *t*(2470) = 1.89, *p* = .058)). On the most general level, linking the L2 after-effect to increased activation in the Multiple Demand system, supports models that posit domain-general bilingual language control (Abutalebi & Green, 2016; Green & Abutalebi, 2013; Wu et al., 2019).

The Multiple Demand network has been claimed to support cognitive control by integrating different types of information and binding them together to support the current cognitive operation (Assem et al., 2020, 2022; Cole et al., 2013; Koechlin & Summerfield, 2007). Activity within the Multiple Demand network has been linked to mechanisms responsible for the implementation and reconfiguration of task sets (Dosenbach et al., 2006, 2007), voluntary attentional control (goal-directed attention; Asplund et al., 2010; Corbetta & Shulman, 2002), and reorienting attention to novel tasks (Corbetta et al., 2008). An important feature of the Multiple Demand network is that it is more engaged in less-automatized and untrained tasks (Shashidhara et al., 2019), thus implying that it implements a domain-general mechanism that supports any difficult or not sufficiently automatized activity that requires additional attentional support. Seen from this perspective, speaking in L1 after L2 may recruit additional domain-general resources to support the execution of a language task. However, this leaves open the question of why exactly speaking in L1 after L2 is so much more difficult than speaking in L1 after L1 that it requires additional support from the Multiple Demand network.

### The L2 after-effect reflects neither word retrieval difficulties nor increased interference between language-specific representations

The L2 after-effect has been previously explained within different theoretical frameworks (see Introduction). While these accounts provide plausible explanations of this effect, no univocal evidence in favor or against these interpretations has been provided (Casado et al., 2022; Wodniecka, Szewczyk, et al., 2020; Declerck & Koch, 2022). All available accounts of the L2 after-effect build on a common theoretical assumption that the difficulty in producing words in L1 after using L2 is a consequence of a control mechanism that changes the balance between the relative activation of L1 and L2 representations (Casado et al., 2022; for a discussion see Declerck & Koch, 2022). Importantly, according to these accounts the change in L1-L2 balance results in increased interference between the two languages that persists for some time after the speaker switches back to speaking in L1 (Casado et al., 2022), which, in turn, increases the difficulty of access and selection of (a) lexico-semantic representations (e.g., Casado et al., 2022) and/or (b) phonological representations and articulatory programs of L1 (e.g., Guo et al., 2011).

The results of the current study do not provide evidence supporting the notion that the L2 after-effect is related to increased interference at the lexico-semantic, phonological, or articulatory levels. We found that the L2 after-effect was not linked to an increased BOLD response within the core language network. Since lexical access and selection mechanisms pertain to core language computations (Hu et al., 2022), difficulties in lexical access should be reflected by an increased BOLD response within the language network. This was, however, not the case in the current study. What is more, we did not find increased brain response to speech production in the L1 after L2 condition in either of the functional networks that were identified using verbal fluency and articulation tasks. Even though both localizer tasks produced robust responses and allowed us to identify a number of functional ROIs, they did not overlap with the neural correlates of the L2 after-effect. Furthermore, we did not find any overlap between the neural correlates of the L2 after-effect and the functional network that was identified using the Stroop task; thus, we found no evidence for increased interference between language representations during speech production in L1 after L2.

The results of the localizer-based analyses provide a rather unexpected characterization of the neural correlates of the L2 after-effect. Contrary to the previous interpretations (Branzi et al., 2014, 2016; Guo et al., 2011; Misra et al., 2012; Declerck & Koch, 2022), the current results indicate that the L2 after-effect reflects neither lexical access difficulty, nor articulatory difficulty, nor the additional engagement of language-specific mechanisms that support word retrieval. In other words, our results suggest that the L2 after-effect does not arise from within the language system itself (or from within language-specific representations). So, how can we then account for the increased difficulty in speaking in the native language after using L2? In the following paragraphs, we propose two putative explanations of the difficulty in production in L1 following the use of L2.

### Putative explanations of the L2 after-effect

#### The L2 after-effect as interference between task schemas

As described in the Introduction, most of the evidence for the involvement of cognitive control in bilingual processing comes from the language-switching paradigm and, more specifically, from the asymmetrical language switch cost (i.e., the finding that the cost of switching to the dominant (L1) language is bigger than to the weaker (L2) language) and language mixing cost effects (i.e., the stronger language (L1) suffers more when both languages are mixed within one block of naming) (Costa & Santesteban, 2004; Gollan & Ferreira, 2009; Meuter & Allport, 1999; Verhoef et al., 2009; for review see: Bobb & Wodniecka, 2013; Declerck et al., 2020; Declerck & Koch, 2022; Declerck & Philipp, 2015). Perhaps the most classic explanation of these effects assumes that they result from the proactive language control that is recruited whenever participants have to name a picture in the weaker L2. This control is recruited to “protect” the weaker L2 (less activated or less automatized) from interference with the stronger L1, either by inhibiting L1 (e.g., Green, 1998; Green & Abutalebi, 2013) or by bolstering the activation of L2 (e.g., Blanco-Elloreta & Caramazza, 2021). When, after using L2, a bilingual speaker switches to her L1, naming in L1 is hampered by the consequences of prior control mechanisms: L1 is now either inhibited, or there is an increased interference with L2, which is still strongly activated.

Explanations that assume the involvement of proactive control have been previously used to account for the L2 after-effect (Wodniecka et al., 2020; Declerck & Koch, 2022), and our current findings are generally consistent with this framework. However, they also point to an important caveat regarding the level of the targeted representations. While models of bilingual language processing typically assume that proactive control targets language-specific representations, such as specific lexical units (see, e.g., BIA+ model, Dijkstra & van Heuven 2002 for bilingual language comprehension; Blanco-Elorrieta & Caramazza, 2021 for bilingual language production), the findings of the current study are incompatible with such views. If proactive control operated on language-specific representations, we should see modulations of brain activity in the after-L1 vs after-L2 conditions in regions directly operating on these language-specific representations, namely in the language network, which was not the case.

Nevertheless, our results support accounts that assume proactive control if one assumes that the control does not target language-specific representations but rather more-abstract ones. Previous literature claims the existence of such representations and refers to them as control representations (Braver et al., 2021) or hierarchical task-schemas (Badre & Nee, 2018). Such representations could be conceived as sets of rules for orchestrating access to language-specific representations – the program we execute when we intend to speak in a given language and which guides all subprocesses involved in subsequent episodes of language production. This may perhaps be best understood in light of the cascade model of executive control (Koechlin & Summerfield, 2007), which model proposes a hierarchy of increasingly abstract control processes. At the sensorimotor control level, a stimulus is simply associated with a reaction (e.g., picking up the phone when it rings). At the contextual control level, the stimulus-response link depends on the current context (not picking up the phone when it rings at your friend’s home). Episodic and branching control levels enable exceptions to the rules under which the current contextual rules can be temporarily and conditionally withheld or reinstated (not picking up the phone at a friend’s place unless he asked you to). Koechlin and Summerfield note that all dual-task situations require such high-level reconfigurations of rules (i.e., task 2 requires temporarily withholding the rulesets of task 1).

Thus, when conceived from the perspective of this framework, the interference between L1 and L2 can occur at the level of rulesets that are managed by these more-abstract levels of cognitive control. The interference occurs between the sets of task rules for speaking in L1 and the sets of task rules for speaking in L2, both of which compete for selection to guide language production. If the interference indeed occurs at the language task-schema level, it would explain why language representations, supported and processed by the language network, are not affected by a prior use of L2, yet it usually slows down the naming process (Branzi et al., 2014; Wodniecka et al., 2020, Casado et al., 2022; Degani et al., 2019). We therefore propose that the activation within the Multiple Demand network may reflect the costs of resolving the interference between L1 and L2 production task schemas.

This idea is in line with the Inhibitory Control Hypothesis (Green, 1998), which proposed that language-switching effects are a consequence of competition between language schemas. However, the Inhibitory Control Hypothesis did not clearly define “language schemas”: they are sometimes described at a more abstract level as sets of rules that guide speech production in different languages, similar to what we propose here; and at other times, they are framed as “regulating the outputs from the lexico-semantic system by altering the activation levels of representations within that system and by inhibiting outputs from that system” (Green, 1998), which implies that they engage in resolution of interference between language-specific representations.

Our proposal is also relevant to a distinction made in the literature regarding bilingual cognitive control, which differentiates between local and global language control. According to this distinction, bilingual language control mechanisms (for example, inhibition) can target not only specific lexical representations (local control) but also the entire language system (global control: Branzi et al., 2014; Van Assche et al., 2013; Wodniecka et al., 2020). The idea that difficulty in bilingual speech production can be driven by increased interference between sets of rules that guide speech production in a given language aligns with the idea of global language control as mechanisms that operate not on language-specific representations (e.g., lemmas, c.f. Van Assche et al., 2013; or nodes linked to a common language node; Dijkstra & van Heuven, 2001; Blanco-Elorrieta & Caramazza, 2021) but on sets of schemas for speaking in a given language.

In sum, while our first tentative explanation of the L2 after-effect builds on a mechanism that has been proposed before (proactive interference during L2 naming), it also points to a critical difference. The key novelty of the proposed interpretation is that the affected representations are not language-specific (lexical units, morphosyntactic units, phonological representations, motor programs) but more general and abstract task-schemas that determine the general programs of using L1 or L2.

#### The L2-after effect as sustained activation of cognitive control

Another possibility that, to the best of our knowledge, has not been proposed in the bilingual language processing literature attributes the after-effects in blocked designs entirely to cognitive control mechanisms without involving any representations at the level of either language or task-schema. This proposal is similar to the first explanation, because it also assumes that a prior speech production task in L2 requires intensified proactive cognitive control to resolve the interference between stronger L1 and weaker L2. However, unlike the first explanation, here we propose that the prior use of L2 leads to increased activation of cognitive control without ever impacting language-specific representations. Instead, the difficult task of speaking in L2 engages cognitive control so much that the control system remains alert even after the task in L2 concludes. We propose that this increased alertness might be achieved by increased engagement of the performance or error-monitoring systems in the brain. As such, the execution of any task (even a non-linguistic one) would proceed in a more controlled way leading to an increase in reaction times. Support for the explanation assuming that the L2 after-effect reflects persisting increased engagement of cognitive control may be found in the Dual Mechanisms of Control framework (Braver, 2021). According to this framework, proactive control recruits the lateral prefrontal cortex, which, unlike the anterior cingulate cortex, is sensitive not only to errors or conflicts in the previous trial but also to control demands specific to the entire task, or the current communicative context (Blanco-Elorrieta & Caramazza, 2021, Casado et al., 2022). Since we did not observe modulations between the L1-after and L2-after conditions in the anterior cingulate cortex activity (see *Supplementary Materials*) but in the lateral prefrontal cortex, which is part of the Multiple Demand system (by-ROI results in the Multiple Demand network are available in *Supplementary Materials*), it can be argued that the L2 after-effect is a manifestation of proactive control.

In sum, the second tentative explanation of the increased brain response related to the L2 after-effect proposes that it reflects the increased engagement of cognitive control triggered by the previous use of L2 and that this engagement lingers during a subsequent task in L1. However, this hypothesis has never been directly tested and, as such, further experiments are needed to falsify it.

While the increased effort related to the L2 after-effect observed within the Multiple Demand network can be putatively explained by increased engagement of proactive mechanisms engaged in performance or error-monitoring systems, other explanations of the observed effect cannot be ruled out. One alternative explanation of the increased engagement of the Multiple Demand network is related to the specificity of the task used in the current study. We used a picture naming task, which requires producing single words. Previous studies on monolingual speakers have shown that the Multiple Demand network responds more strongly to naming objects (i.e., producing isolated words) than to describing events presented with pictures (i.e., producing full sentences; Hu et al., 2022). Similar effects have also been found for language comprehension: the Multiple Demand network was shown to respond more strongly to listening to lists of unconnected words than full sentences (Diachek et al., 2020). Altogether these results suggests that the Multiple Demand network does not support the core aspects of language production. This does not rule out the possibility that the engagement of the domain-general network reflects increased engagement of non-linguistic control processes in the production of words in L1 after L2 in contrast to L1 after L1; however, it may suggest that the L2 after-effect measured in a picture naming task may be “inflated” by the mere demands related to producing single, isolated words. As such, the effect might reflect a laboratory phenomenon which not necessarily exemplifies cognitive control mechanisms that bilinguals engage in more natural conversations of everyday language use. As such, further studies implementing more naturalistic paradigms that require bilinguals to produce more complex outputs are needed to separate the components driven by task-related demands from the actual engagement of cognitive control in the L2 after-effect.

## Conclusions

The novelty of our contribution is twofold: First, to identify the cognitive mechanisms that correspond to the neural signature of the L2 after-effect, we used the functional localizer approach (Fedorenko et al., 2010), which to the best of our knowledge has never been used in experiments looking at the brain basis of bilingual speech production. Second, unlike previous studies, our conclusions are based on a design that was optimized to measure the index of language control, i.e., the L2 after-effect, while reducing many possible confounds. We believe that the combination of precise analytical tools, carefully designed tasks, as well as the relatively large number of participants tested (n=42) lead to new insights into the neurocognitive mechanisms of language control in bilinguals.

In contrast to the dominating theories and views in the field, we found that the L2 after-effect is unlikely to arise as a result of interference between language-specific representations in L1 and L2 because it does not overlap with the language network, or with any task-specific localizer that targets specific language processes (lexical access, articulation, language interference, and inhibition). Instead, we found that the L2 after-effect overlaps with the Multiple Demand network and thus reflects a domain-general mechanism. We propose that the L2 after-effect may manifest as either increased interference between high-level task schemas used to orchestrate language use in L1 and L2, or as a lingering increase in the engagement of domain-general cognitive control mechanisms in speech production.

## Data/code availability statement

The raw neuroimaging data collected for this study along with the data on participants’ language experience is available at the OpenNeuro repository https://openneuro.org/datasets/ds004456.

Data and code necessary to reproduce the presented ROI and behavioral analyses is available at the https://osf.io/59za8/?view_only=e23f3500d12e4109b75ca18fdbd320e5.

## Acknowledgements

The authors would like to thank all members of our Psychology of Language and Bilingualism Laboratory LangUsta who contributed to the research project by collecting and coding the data, all participants who took part in the study, as well as Mike Timberlake who proofread the text. We are very grateful to Evelina Fedorenko for inspiration and insightful comments on the design and results.

