## Supplementary Materials for "Tracking components of bilingual language control in speech production: an fMRI study using functional localizers"

### By-ROI analyses for each functional network

To further explore the fine-grained differences between our critical conditions that might only affect a subset of functional parcels within each network, we fitted separate models for each ROI in each of the functional networks. To this aim, we fitted a linear mixed-effect model for each ROI using the following formula:

% signal change ~ PrecedingLanguage +

(1 + PrecedingLanguage | Subject)

The results corresponding to differences between speaking in *L1 after L2* and *L1 after L1* for each ROI within the five functional networks are presented in Tables S1-S5 and Figures S1-S5.

#### Summary of the results

This appendix presents analyses of the *L2 after-effect,* i.e. the difference in brain response to speaking in *L1 after L2* and *L1 after L1,* carried out for each ROI within the Multiple Demand and Language networks as well as within the functional networks identified with an articulation, Verbal Fluency, and Stroop tasks. No significant differences were found for neither of the ROIs within the Language network as well as any of the three task-specific networks: Articulation, Verbal Fluency, and Stroop. Within the Multiple Demand network, significant differences between brain response estimates to speaking in *L1 after L2* and *L1 after L1* were found in the left and right parietal lobules (angular gyrus extending to the superior parietal lobule), right middle frontal gyrus as well as in the right frontal pole. Previous studies proposed that the right and left inferior parietal lobules (IPLs), where the angular gyrus is located, are the brain basis of attentional adjustments of bilingual language control, with the left IPL being responsible for biasing language selection away from the language not in use and the right IPL biasing the attention towards the language in use (Abutalebi & Green, 2016; Calabria et al., 2018; Wu et al., 2019). Moreover, brain activity corresponding to mechanisms of sustained language control has been linked to increased response in the right middle frontal (Collette et al., 2005; Wang et al., 2009) and prefrontal cortices (Braver et al., 2003; Wang et al., 2009). Bilateral parietal lobules as well as the right MFG and frontal pole correspond to the group-level functional ROIs within the MD network that showed significant response in our study. As such, the results of a by-ROI analysis further support the conclusions drawn based on the network-level analyses: that the *L2 after-effect* reflects the increased engagement of sustained, proactive control mechanisms.

Four functional ROIs within the MD network showed significant response to the L1 after L2 > L1 after L1 contrast. Significant response to this contrast was found in the functional ROIs spanning the left and right Angular Gyrus, right SFG/MFG and right Frontal Pole. Additionally, given substantial effect size estimates for all these ROIs (*d* > 0.6), we believe these results may provide useful guides for future research characterizing the Multiple Demand network as well as research focused on bilingual language control and the *L2 after-effect.* Below we provide a short characterization of the functional profile of the ROIs in which we found a significant *L2 after-effect.*

#### Multiple Demand network

**Table S1.** **Results of linear mixed-effect models fitted for each functional ROI within the MD network.** Presented estimates correspond to the effect of the preceding language (*L1 after L1* vs *L1 after L2*). Anatomical labels were derived from Harvard-Oxford Cortical, Harvard-Oxford Subcortical, or Cereberall probabilitistic atlases (from FSL) and they correspond to one or two labels with the highest probabilistic overlap with each functional ROI.

| **Multiple Demand network** | | | | | | | |
| --- | --- | --- | --- | --- | --- | --- | --- |
| **Region of interest** | | **Fixed effect of condition**  *(L1 after L2 > L1 after L1)* | | | | | |
| **hemisphere** | **label** | **Estimate** | **Standard  Error** | ***t* - value** | **Effect size *d*** | ***p*-value** | ***p-*value FDR corrected** |
| LH | Lateral Occipital Cortex, superior division | 0,09 | 0,05 | 1,58 | 0,50 | 0,121 | 0,221 |
|  | Precuneous Cortex |  |  |  |  |  |  |
| LH | Angular Gyrus | 0,17 | 0,06 | 2,84 | 0,90 | 0,007 | 0,041* |
|  | Superior Parietal Lobule |  |  |  |  |  |  |
| LH | Supramarginal Gyrus, anterior division | 0,02 | 0,03 | 0,67 | 0,21 | 0,510 | 0,598 |
|  | Postcentral Gyrus |  |  |  |  |  |  |
| LH | Superior Frontal Gyrus | 0,10 | 0,05 | 2,14 | 0,68 | 0,038 | 0,099 |
|  | Middle Frontal Gyrus |  |  |  |  |  |  |
| LH | Precentral Gyrus | 0,05 | 0,05 | 0,97 | 0,31 | 0,338 | 0,450 |
|  | Middle Frontal Gyrus |  |  |  |  |  |  |
| LH | Inferior Frontal Gyrus, pars opercularis | 0,02 | 0,05 | 0,35 | 0,11 | 0,732 | 0,734 |
| LH | Middle Frontal Gyrus | 0,09 | 0,04 | 2,13 | 0,67 | 0,040 | 0,099 |
| LH | Frontal Pole | 0,10 | 0,05 | 2,06 | 0,65 | 0,046 | 0,101 |
| LH | Insular Cortex | -0,01 | 0,03 | -0,34 | -0,11 | 0,734 | 0,734 |
| LH | Paracingulate Gyrus | 0,04 | 0,04 | 1,24 | 0,39 | 0,223 | 0,343 |
| RH | Lateral Occipital Cortex, superior division | 0,10 | 0,06 | 1,77 | 0,56 | 0,084 | 0,169 |
|  | Precuneous Cortex |  |  |  |  |  |  |
| RH | Angular Gyrus | 0,16 | 0,05 | 3,05 | 0,97 | 0,004 | 0,041* |
|  | Superior Parietal Lobule |  |  |  |  |  |  |
| RH | Supramarginal Gyrus, anterior division | 0,08 | 0,03 | 2,46 | 0,78 | 0,018 | 0,060 |
|  | Postcentral Gyrus |  |  |  |  |  |  |
| RH | Superior Frontal Gyrus | 0,10 | 0,04 | 2,79 | 0,88 | 0,008 | 0,041* |
|  | Middle Frontal Gyrus |  |  |  |  |  |  |
| RH | Precentral Gyrus | 0,05 | 0,04 | 1,08 | 0,34 | 0,285 | 0,407 |
|  | Middle Frontal Gyrus |  |  |  |  |  |  |
| RH | Inferior Frontal Gyrus, pars opercularis | 0,03 | 0,03 | 0,90 | 0,29 | 0,373 | 0,466 |
| RH | Middle Frontal Gyrus | 0,10 | 0,04 | 2,54 | 0,80 | 0,015 | 0,060 |
| RH | Frontal Pole | 0,17 | 0,06 | 3,01 | 0,95 | 0,005 | 0,041* |
| RH | Insular Cortex | -0,05 | 0,03 | -1,41 | -0,45 | 0,167 | 0,278 |
| RH | Paracingulate Gyrus | 0,02 | 0,03 | 0,62 | 0,20 | 0,538 | 0,598 |

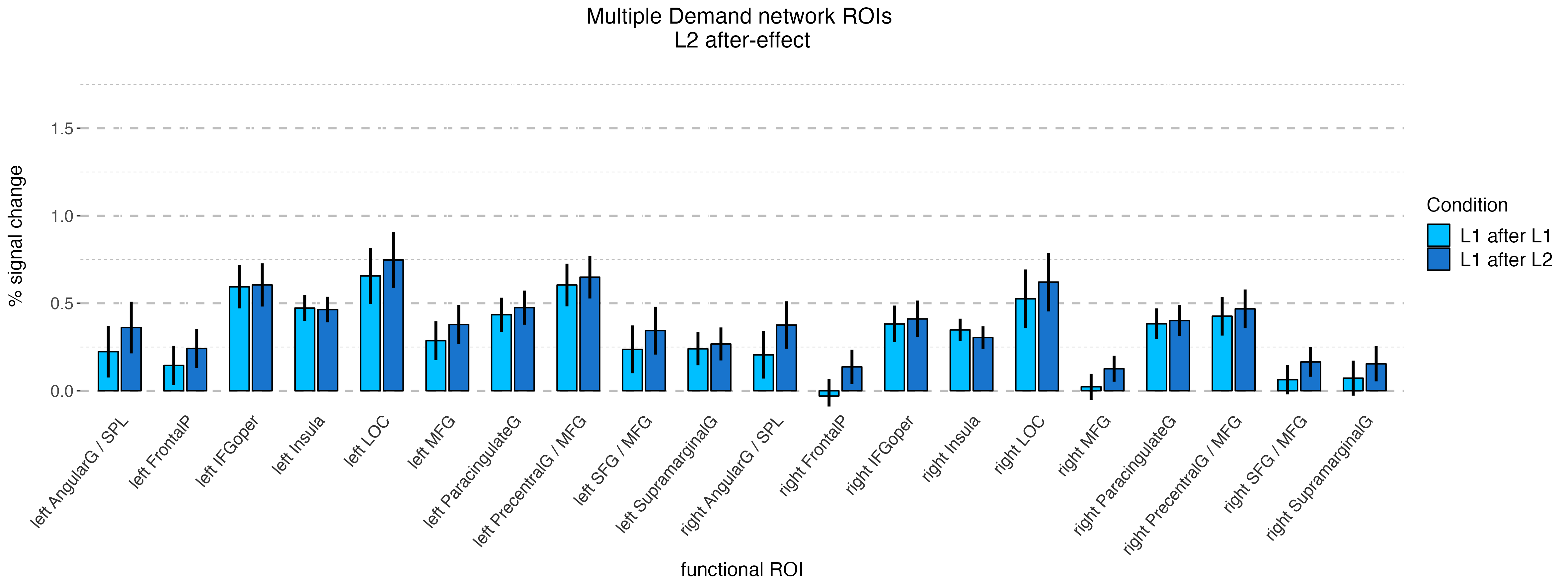
**Figure S1.** **Mean percent signal change in each functional ROI of the Multiple Demand network.** Errorbars represent low and high confidence interval for the effect of condition (L1 after L1 vs L1 after L2) in each fROI.

#### Language network

**Table S2.** **Results of linear mixed-effect models fitted for each functional ROI within the Language network.** Presented estimates correspond to the effect of the preceding language (*L1 after L1* vs *L1 after L2*). Anatomical labels were derived from Harvard-Oxford Cortical, Harvard-Oxford Subcortical, or Cereberall probabilitistic atlases (from FSL) and they correspond to one or two labels with the highest probabilistic overlap with each functional ROI.

| **Language network** | | | | | | | |
| --- | --- | --- | --- | --- | --- | --- | --- |
| **Region of interest** | | **Fixed effect of condition**  *(L1 after L2 > L1 after L1)* | | | | | |
| **hemisphere** | **label** | **Estimate** | **Standard  Error** | ***t* - value** | **Effect size *d*** | ***p*-value** | ***p-*value FDR corrected** |
| LH | Frontal Orbital Cortex | 0,00 | 0,05 | -0,02 | 0,00 | 0,988 | 0,988 |
|  | Inferior Frontal Gyrus, pars triangularis |  |  |  |  |  |  |
| LH | Inferior Frontal Gyrus, pars opercularis | 0,01 | 0,04 | 0,29 | 0,09 | 0,776 | 0,944 |
| LH | Precentral Gyrus | 0,05 | 0,09 | 0,61 | 0,19 | 0,545 | 0,885 |
|  | Middle Frontal Gyrus |  |  |  |  |  |  |
| LH | Temporal Pole | 0,02 | 0,03 | 0,67 | 0,21 | 0,509 | 0,885 |
| LH | Angular Gyrus | 0,05 | 0,03 | 1,46 | 0,46 | 0,152 | 0,885 |
|  | Middle Temporal Gyrus, temporooccipital part |  |  |  |  |  |  |
| LH | Lateral Occipital Cortex, superior division | 0,01 | 0,05 | 0,13 | 0,04 | 0,900 | 0,982 |
| RH | Frontal Orbital Cortex | 0,01 | 0,05 | 0,27 | 0,09 | 0,787 | 0,944 |
|  | Inferior Frontal Gyrus, pars triangularis |  |  |  |  |  |  |
| RH | Inferior Frontal Gyrus, pars opercularis | 0,03 | 0,04 | 0,59 | 0,19 | 0,559 | 0,885 |
| RH | Precentral Gyrus | 0,05 | 0,08 | 0,69 | 0,22 | 0,495 | 0,885 |
|  | Middle Frontal Gyrus |  |  |  |  |  |  |
| RH | Temporal Pole | 0,02 | 0,04 | 0,54 | 0,17 | 0,590 | 0,885 |
| RH | Angular Gyrus | 0,03 | 0,04 | 0,70 | 0,22 | 0,486 | 0,885 |
|  | Middle Temporal Gyrus, temporooccipital part |  |  |  |  |  |  |
| RH | Lateral Occipital Cortex, superior division | 0,03 | 0,05 | 0,63 | 0,20 | 0,534 | 0,885 |

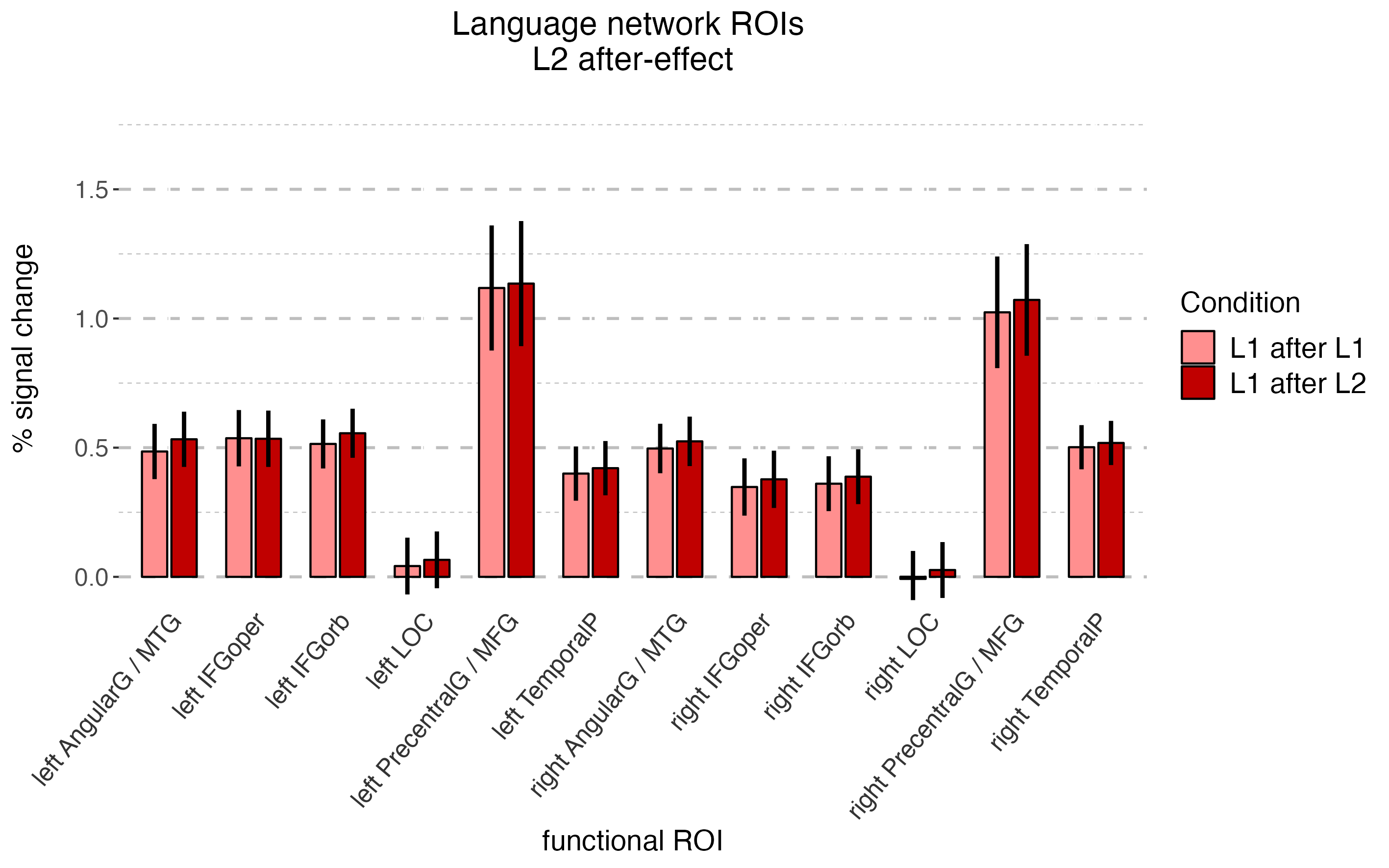

**Figure S2.** **Mean percent signal change in each functional ROI of the Language network.** Errorbars represent low and high confidence interval for the effect of condition (L1 after L1 vs L1 after L2) in each fROI.

#### Articulation network

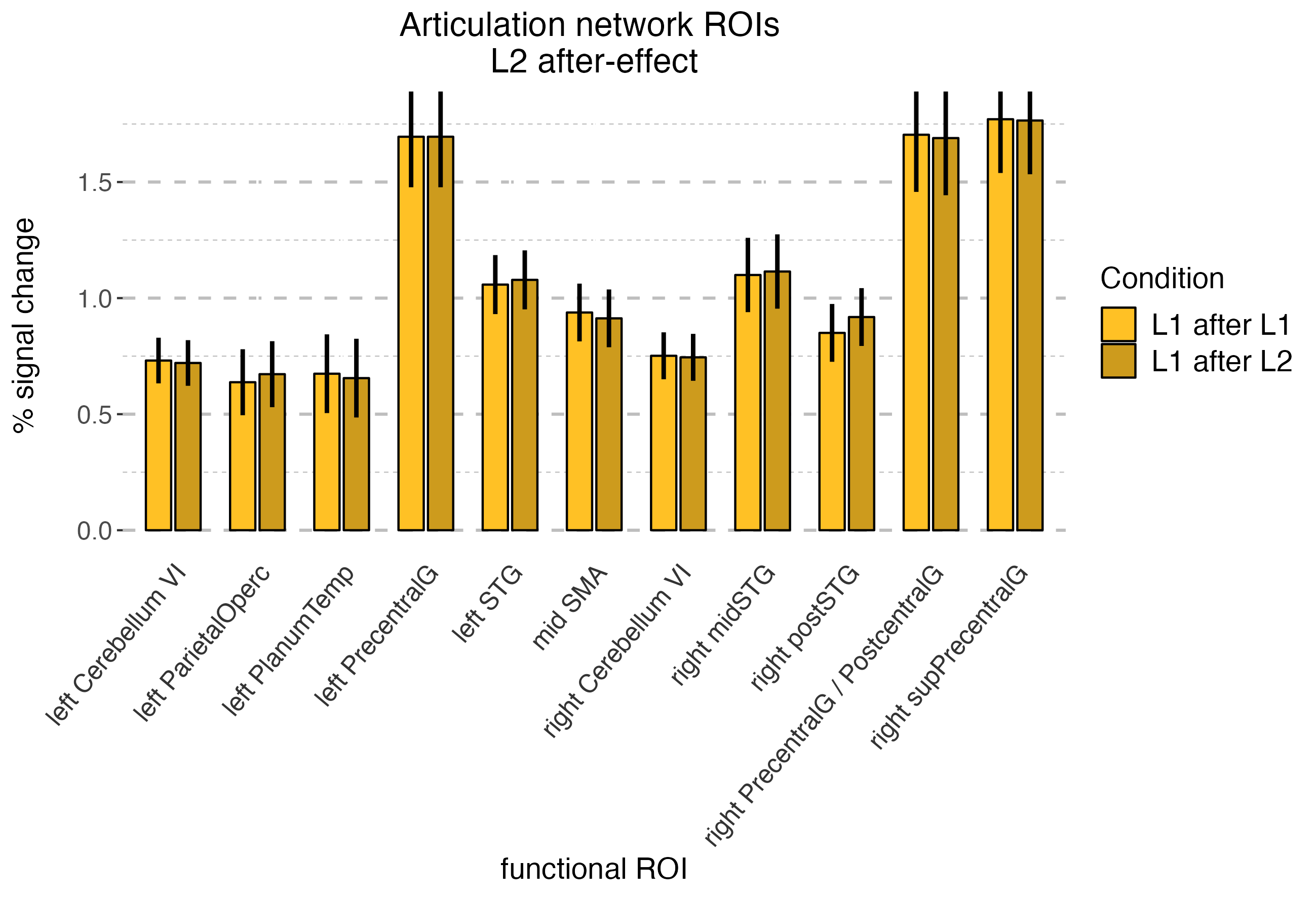
**Table S3.** **Results of linear mixed-effect models fitted for each functional ROI within the Articulation network.** Presented estimates correspond to the effect of the preceding language (*L1 after L1* vs *L1 after L2*). Anatomical labels were derived from Harvard-Oxford Cortical, Harvard-Oxford Subcortical, or Cereberall probabilitistic atlases (from FSL) and they correspond to one or two labels with the highest probabilistic overlap with each functional ROI.

| **Articulation network** | | | | | | | | | | | | | | |
| --- | --- | --- | --- | --- | --- | --- | --- | --- | --- | --- | --- | --- | --- | --- |
| **Region of interest** | | **Fixed effect of condition**  *(L1 after L2 > L1 after L1)* | | | | | | | | | | | | |
| **hemisphere** | **label** | | | **Estimate** | **Standard  Error** | | ***t* - value** | | **Effect size *d*** | | ***p*-value** | | ***p-*value FDR corrected** | |
| LH | Superior Temporal Gyrus, posterior division | | 0,02 | | | 0,07 | | 0,31 | | 0,10 | | 0,760 | | 0,999 |
| LH | Parietal Operculum Cortex | | 0,03 | | | 0,05 | | 0,69 | | 0,22 | | 0,498 | | 0,999 |
| LH | Planum Temporale | | -0,02 | | | 0,07 | | -0,26 | | -0,08 | | 0,800 | | 0,999 |
| LH | Precentral Gyrus | | 0,00 | | | 0,09 | | 0,00 | | 0,00 | | 0,999 | | 0,999 |
| RH | Superior Temporal Gyrus, posterior division | | 0,01 | | | 0,07 | | 0,21 | | 0,07 | | 0,835 | | 0,999 |
| RH | Superior Temporal Gyrus, posterior division | | 0,07 | | | 0,05 | | 1,39 | | 0,44 | | 0,172 | | 0,999 |
| RH | Precentral Gyrus | | -0,01 | | | 0,12 | | -0,13 | | -0,04 | | 0,901 | | 0,999 |
|  | Postcentral Gyrus | |  |  |  |  |  |  |  |  |  |  |  |  |
| RH | Precentral Gyrus | | -0,01 | | | 0,10 | | -0,05 | | -0,02 | | 0,957 | | 0,999 |
| Mid | Supplementary Motor Cortex | | -0,03 | | | 0,05 | | -0,52 | | -0,16 | | 0,607 | | 0,999 |
| RH | Cerebellum Right VI | | -0,01 | | | 0,03 | | -0,20 | | -0,06 | | 0,845 | | 0,999 |
| LH | Cerebellum Left VI | | -0,01 | | | 0,03 | | -0,32 | | -0,10 | | 0,753 | | 0,999 |

**Figure S3.** **Mean percent signal change in each functional ROI of the Articulation network.** Errorbars represent low and high confidence interval for the effect of condition (L1 after L1 vs L1 after L2) in each fROI.

#### Verbal Fluency network

**Table S4.** **Results of linear mixed-effect models fitted for each functional ROI within the Articulation network.** Presented estimates correspond to the effect of the preceding language (*L1 after L1* vs *L1 after L2*). Anatomical labels were derived from Harvard-Oxford Cortical, Harvard-Oxford Subcortical, or Cereberall probabilitistic atlases (from FSL) and they correspond to one or two labels with the highest probabilistic overlap with each functional ROI.

| **Verbal Fluency network** | | | | | | | |
| --- | --- | --- | --- | --- | --- | --- | --- |
| **Region of interest** | | **Fixed effect of condition**  *(L1 after L2 > L1 after L1)* | | | | | |
| **hemisphere** | **label** | **Estimate** | **Standard  Error** | ***t* - value** | **Effect size *d*** | ***p*-value** | ***p-*value FDR corrected** |
| LH | Middle Frontal Gyrus | -0,01 | 0,05 | -0,16 | -0,05 | 0,871 | 0,871 |
|  | Inferior Frontal Gyrus, pars opercularis |  |  |  |  |  |  |
| LH | Frontal Pole | -0,02 | 0,04 | -0,58 | -0,18 | 0,565 | 0,848 |
|  | Inferior Frontal Gyrus, pars triangularis |  |  |  |  |  |  |
| LH | Paracingulate Gyrus | 0,05 | 0,04 | 1,21 | 0,38 | 0,232 | 0,768 |
|  | Superior Frontal Gyrus |  |  |  |  |  |  |
| RH | Cerebellum Right Crus II | -0,06 | 0,10 | -0,62 | -0,20 | 0,538 | 0,848 |
| bilateral | Occipital Pole | -0,03 | 0,09 | -0,36 | -0,12 | 0,718 | 0,848 |
| RH | Cerebellum Right Crus I | -0,02 | 0,08 | -0,21 | -0,07 | 0,833 | 0,871 |
| LH | Insular Cortex | -0,01 | 0,04 | -0,37 | -0,12 | 0,716 | 0,848 |
|  | Frontal Orbital Cortex |  |  |  |  |  |  |
| LH | Frontal Orbital Cortex | 0,08 | 0,05 | 1,49 | 0,47 | 0,144 | 0,768 |
| LH | Middle Frontal Gyrus | 0,07 | 0,06 | 1,30 | 0,41 | 0,202 | 0,768 |
|  | Superior Frontal Gyrus |  |  |  |  |  |  |
| LH | Inferior Temporal Gyrus, temporooccipital part | -0,03 | 0,05 | -0,56 | -0,18 | 0,580 | 0,848 |
| RH | Cerebellum Right Crus II | 0,02 | 0,05 | 0,43 | 0,14 | 0,669 | 0,848 |
| LH | Lateral Occipital Cortex, superior division | -0,05 | 0,04 | -1,20 | -0,38 | 0,236 | 0,768 |
| LH | Lingual Gyrus | 0,06 | 0,07 | 0,82 | 0,26 | 0,417 | 0,848 |
|  | Precuneous Cortex |  |  |  |  |  |  |

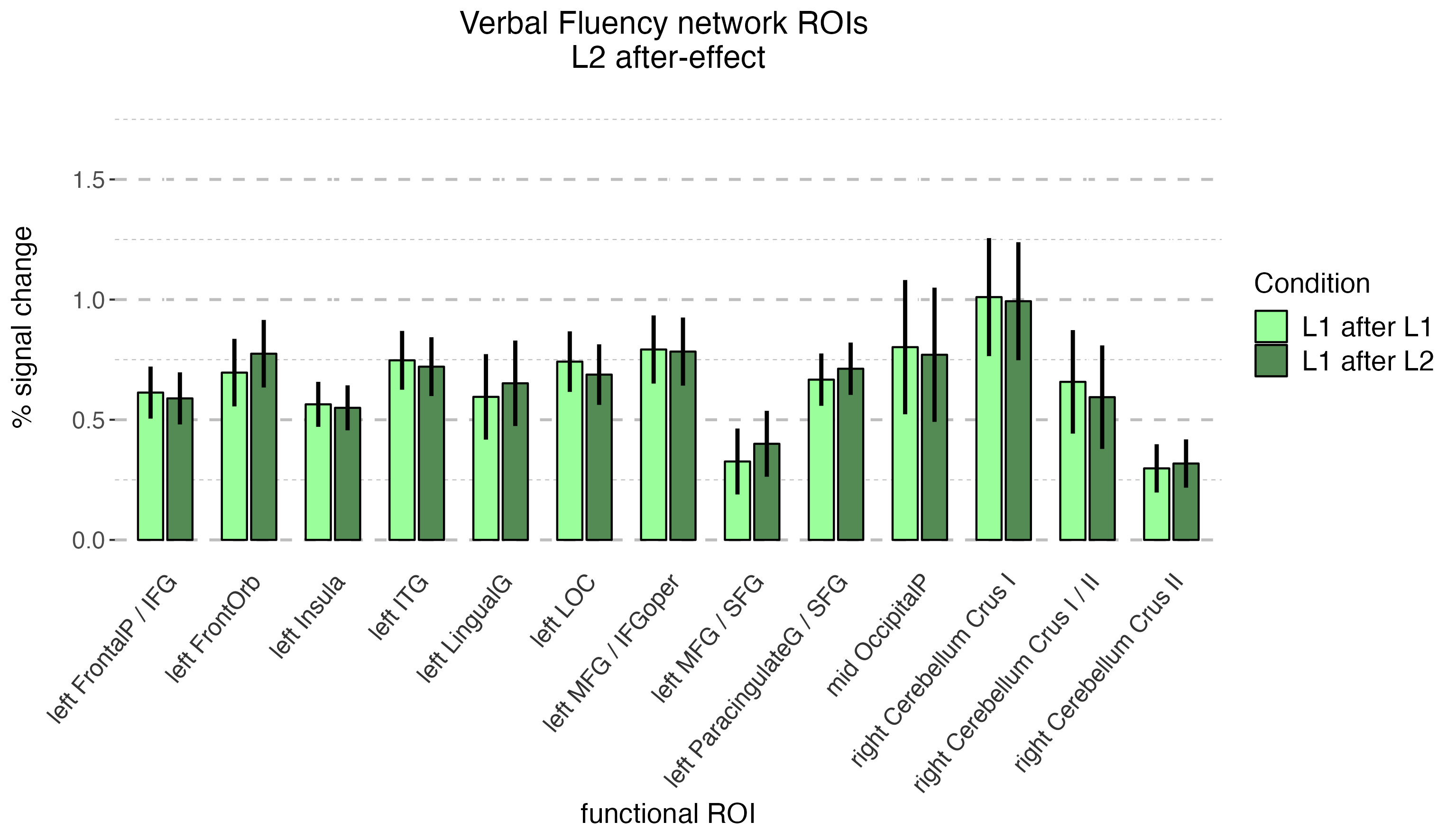
**Figure S4.** **Mean percent signal change in each functional ROI of the Verbal Fluency network.** Errorbars represent low and high confidence interval for the effect of condition (L1 after L1 vs L1 after L2) in each fROI.

#### Stroop network

**Table S5.** **Results of linear mixed-effect models fitted for each functional ROI within the Articulation network.** Presented estimates correspond to the effect of the preceding language (*L1 after L1* vs *L1 after L2*). Anatomical labels were derived from Harvard-Oxford Cortical, Harvard-Oxford Subcortical, or Cereberall probabilitistic atlases (from FSL) and they correspond to one or two labels with the highest probabilistic overlap with each functional ROI.

| **Stroop network** | | | | | | | |
| --- | --- | --- | --- | --- | --- | --- | --- |
| **Region of interest** | | **Fixed effect of condition**  *(L1 after L2 > L1 after L1)* | | | | | |
| **hemisphere** | **label** | **Estimate** | **Standard  Error** | ***t* - value** | **Effect size *d*** | ***p*-value** | ***p-*value FDR corrected** |
| LH | Lateral Occipital Cortex, superior division | 0,02 | 0,05 | 0,33 | 0,10 | 0,744 | 0,969 |
| LH | Supramarginal Gyrus | 0,03 | 0,05 | 0,61 | 0,19 | 0,545 | 0,886 |
| LH | Superior Parietal Lobule | 0,00 | 0,04 | -0,13 | -0,04 | 0,895 | 0,969 |
| LH | Middle Frontal Gyrus | 0,02 | 0,05 | 0,29 | 0,09 | 0,777 | 0,969 |
|  | Inferior Frontal Gyrus, pars opercularis |  |  |  |  |  |  |
| RH | Supramarginal Gyrus, posterior division | 0,04 | 0,04 | 1,22 | 0,39 | 0,229 | 0,693 |
|  | Superior Parietal Lobule |  |  |  |  |  |  |
| LH | Middle Frontal Gyrus | 0,05 | 0,05 | 1,01 | 0,32 | 0,320 | 0,693 |
|  | Frontal Pole |  |  |  |  |  |  |
| RH | Lateral Occipital Cortex, superior division | 0,08 | 0,06 | 1,38 | 0,44 | 0,176 | 0,693 |
| RH | Middle Frontal Gyrus | 0,00 | 0,05 | 0,01 | 0,00 | 0,991 | 0,991 |
|  | Inferior Frontal Gyrus, pars opercularis |  |  |  |  |  |  |
| RH | Middle Frontal Gyrus | 0,08 | 0,05 | 1,43 | 0,45 | 0,160 | 0,693 |
|  | Superior Frontal Gyrus |  |  |  |  |  |  |
| LH | Inferior Temporal Gyrus, temporooccipital part | -0,01 | 0,06 | -0,18 | -0,06 | 0,858 | 0,969 |
| Mid | Paracingulate Gyrus | 0,08 | 0,06 | 1,48 | 0,47 | 0,146 | 0,693 |
| RH | Middle Frontal Gyrus | 0,02 | 0,03 | 0,64 | 0,20 | 0,529 | 0,886 |
|  | Superior Frontal Gyrus |  |  |  |  |  |  |
| RH | Frontal Pole | 0,05 | 0,05 | 1,01 | 0,32 | 0,319 | 0,693 |
|  | Middle Frontal Gyrus |  |  |  |  |  |  |

**Figure S5.** **Mean percent signal change in each functional ROI of the Stroop network.** Errorbars represent low and high confidence interval for the effect of condition (L1 after L1 vs L1 after L2) in each fROI.
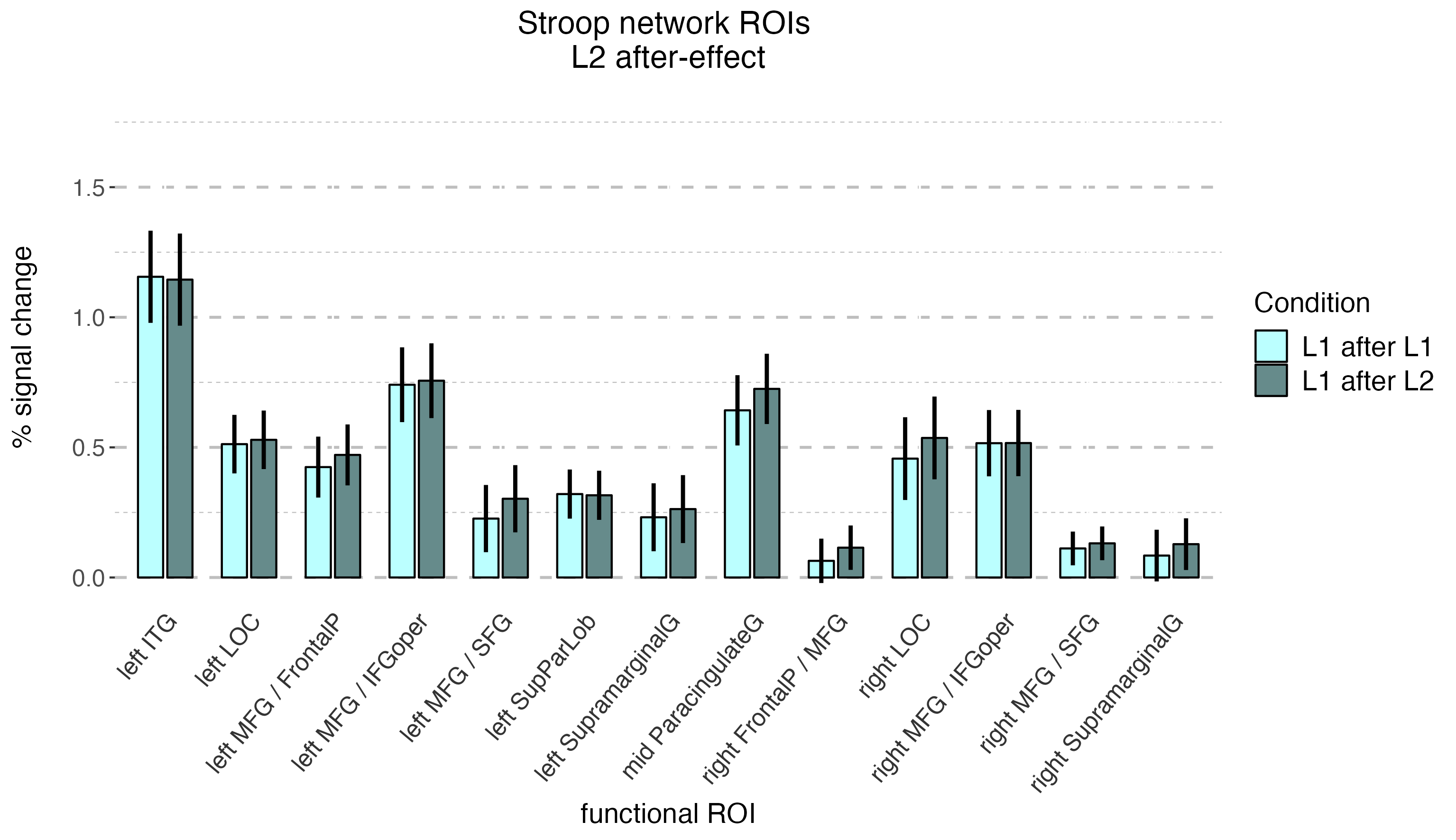

### Characterization of the functional networks: response to localizer conditions within different functional networks

To better charactrerize the functional networks used in this study, we have run additional analyses in which we extracted responses to the tasks used as localizers within the functional networks used to explore the *L2 after-effect.* These analyses provide a functional characterization of the networks identified with the language and Multiple Demand localizerts as well as the Articulation, Stroop, and Verbal Fluency task. These analyses were conducted using the same pipeline as the one used in the localizer-based analyses of the responses to *L1-after-L2* and *L1-after-L1* in the Picture Naming task.

In the first analysis we explored the responses to all five localizer tasks within the langauge and Multiple Demand networks. As the parcels used to for these two networks were symmetrical between hemispheres, in the models we also included the interactions of condition (i.e., conditions of a given localizer task) with the hemisphere, to account for differences in response between the left and right hemisphere fROIs. To this aim, we fitted a linear mixed-effect model using the following formula:

% signal change ~ Conditon + Hemisphere + Condition:Hemisphere +

(1 + Conditon + Hemisphere | Subject) +

(1 | ROI)

Before the analysis, categorical predictors were deviation-coded. The analysis was performed using the lmer() function from the lmerTest package (Kuznetsova et al., 2017). Furthermore, pairwse comparisons were conducted to explore differences between conditions in each hemisphere separately, using the emmeans() function from the emmenas package.

In the second analysis, we explored the responses to the Language and Multiple Demand localizer tasks (i.e., intact vs. degraded speech and hard vs. easy visual working memory) within the language, Multiple Demand, Articulaton, Stroop, and Verbal Fluency networks. As the fROIs for the Articulation, Stroop, and Verbal Fluency networks are not symmetrical, i.e., thy differ in number and location between hemispheres, in this analysis we modelled responses to the Language and Multiple Demand localizers in the entire networks, without differentiating between hemispheres. To this aim, we fitted a linear mixed-effect model using the following formula:

% signal change ~ Conditon +

(1 + Conditon | Subject) +

(1 | ROI)

Similarly to the first analysis, categorical predictor of condition was deviation-coded and the analysis was performed using the lmer() function from the lmerTest package (Kuznetsova et al., 2017).

#### Response to the localizer tasks within the Language and Multiple Demand networks

Within the language network, we found significant differences between the localizer conditions for the langauge localizer task and the Articulation task. Furthermore, in both these tasks we found interactions between condition and hemisphere: in the language localizer task difference in activation between the intact and degraded speech was larger in the left (β = 0.450, *t(68)* = 12.306, *p* < .001) than the right hemisphere (β = 0.235, *t(68)* = 6.413, *p* < .001); on the contrary, in the Articulation task difference in activation between repeating syllables and a motor sequence was larger in the right (β = 0.323, *t(55)* = 6.825, *p* < .001) than the left hemisphere (β = 0.234, *t(55)* = 4.973, *p* < .001). No differences between task conditions were found for the Multiple Demand localizer and for the Stroop task. However, it is important to note that while the visual working memory task, in neither its hard or easy version, yielded responses significantly different than zero, both conditions of the Stroop task, which is a linguistic task requiring speech production, had non-zero responses in the language network. Finally, even though we did not find a main effect of condition in the Verbal Fluency task, there was a significant interaction of condition and hemisphere: in the left hemisphere we found stronger responses to the fluency task compared to the automated speech baseline (β = 0.242, *t(63)* = 5.389, *p* < .001) and the opposite effect in the right hemisphere, with stronger responses to the automated speech condition than the verbal fluency task (β = -0.237, *t(63)* = -5.289, *p* < .001). The results of analyses within the language network are presented in Table S6 and Figure S6.

**Table S6. Results of linear mixed-effect models fitted for each localizer task within the language network.** The table presents effects of condition, hemisphere, and their interaction for the Language localizer, MD localizer, Articulation, Stroop, and Verbal Fluency tasks.

| **Task** | **Effect** | **Estimate** | **Standard  Error** | ***t* - value** | **Effect size *d*** | ***p*-value** |
| --- | --- | --- | --- | --- | --- | --- |
| **Language localizer** | listening to intact speech > degraded speech | -0,34 | 0,03 | -10,70 | -3,38 | 0,000 |
|  | left > right | -0,13 | 0,21 | -0,61 | -0,38 | 0,557 |
|  | interaction | 0,22 | 0,04 | 6,08 | 0,28 | 0,000 |
| **MD localizer** | hard vWM > easy vWM | 0,03 | 0,02 | 1,28 | 0,40 | 0,209 |
|  | left > right | 0,04 | 0,13 | 0,33 | 0,20 | 0,746 |
|  | interaction | -0,01 | 0,03 | -0,45 | -0,02 | 0,651 |
| **Articulation** | syllable repetition > motor sequence | -0,28 | 0,04 | -6,41 | -2,03 | 0,000 |
|  | left > right | 0,00 | 0,14 | -0,01 | 0,00 | 0,995 |
|  | interaction | -0,09 | 0,04 | -2,43 | -0,11 | 0,015 |
| **Stroop** | naming color words > adjectives | -0,02 | 0,02 | -0,82 | -0,07 | 0,414 |
|  | left > right | -0,01 | 0,30 | -0,05 | -0,03 | 0,964 |
|  | interaction | 0,01 | 0,05 | 0,27 | 0,01 | 0,785 |
| **Verbal Fluency** | fluency > automated speech | 0,00 | 0,04 | -0,06 | -0,02 | 0,955 |
|  | left > right | 0,03 | 0,27 | 0,11 | 0,07 | 0,918 |
|  | interaction | 0,48 | 0,04 | 11,89 | 0,56 | 0,000 |

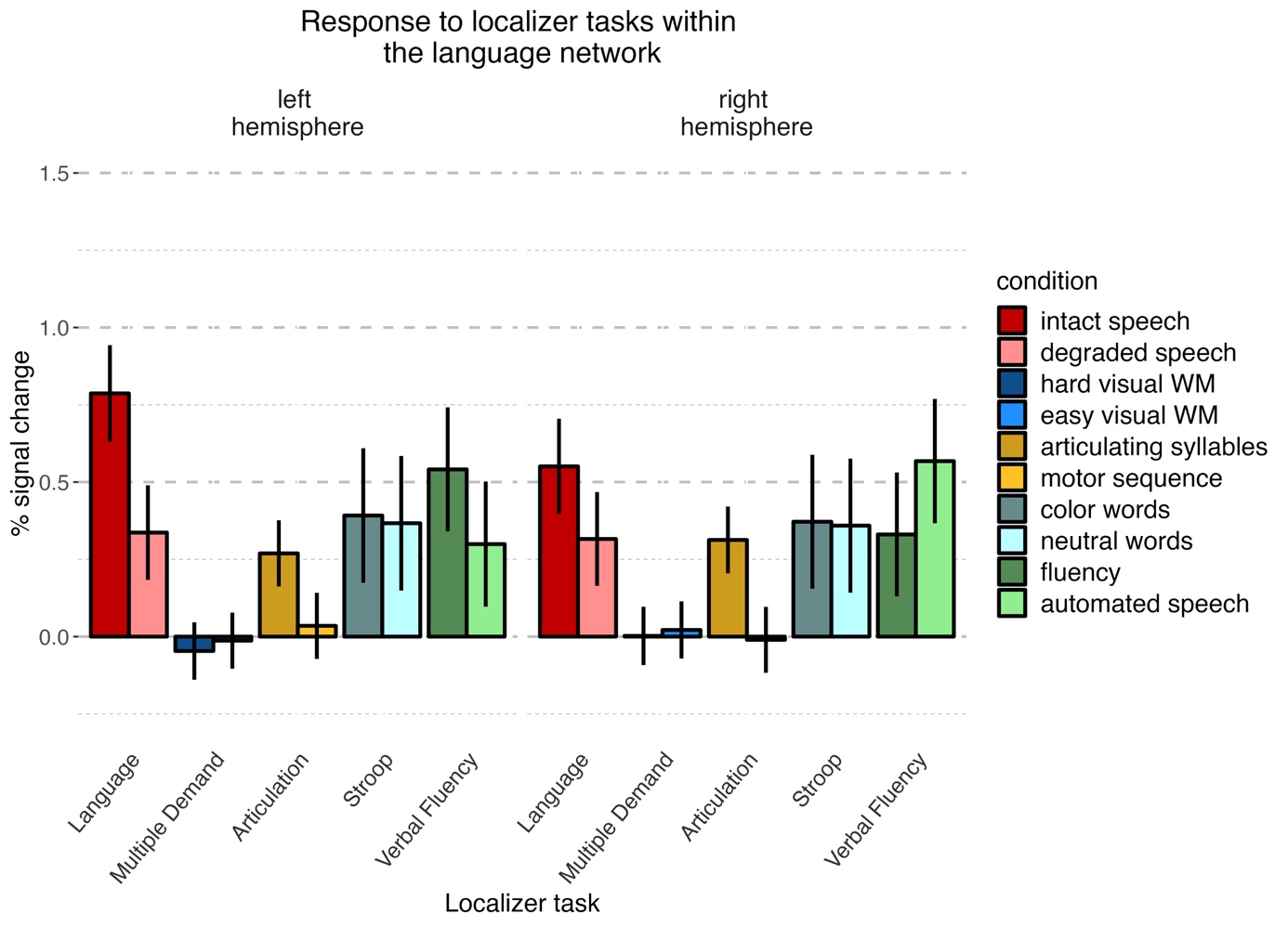

**Figure S6.** **Mean percent signal change corresponding to the localizer task conditions in the language network.** Errorbars represent standard error of the effect of condition in each task and hemisphere.

Within the Multiple Demand network we found significant differences between the localizer conditions for the Language localizer task, the Multiple Demand localizer task, the Articulation task, and for the Stroop task. For the Language localizer task, the effect found in the Multiple Demand network was opposite to the one found in the language network: the MD network responded more strongly to listening to degraded speech than to intact speech. We have also found a significant interaction of condition and hemisphere showing that the difference in activation between degraded and intact speech was larger in the left (β = -0.092, *t(59)* = -3.713, *p* = .001) than in the right hemisphere (β = -0.042, *t(59)* = -1.692, *p* = .183). The opposite effect to the one observed in the language network was also found for the Articulation task: stronger responses were observed for the motor sequence than for repeating syllables. For the Stroop task, stronger responses were observed for the color words compared to the neutral words. A significant interaction with the hemisphere was also found, revealing that the difference between color and neutral words was stronger in the left (β = 0.304, *t(61)* = 8.504, *p* < .001) than in the right hemisphere (β = 0.225, *t(61)* = 6.290, *p* < .001). Finally, even though no main effect of condition was found for the Verbal Fleuncy task, similarly to the language network, an interaction of condition and hemisphere was found: : in the left hemisphere we found stronger responses to the fluency task compared to the automated speech baseline (β = 0.389, *t(45)* = 6.660, *p* < .001) and the opposite effect in the right hemisphere, with stronger responses to the automated speech condition than the verbal fluency task (β = -0.307, *t(45)* = -5.260, *p* < .001). The results of analyses within the language network are presented in Table S7 and Figure S7.

**Table S7. Results of linear mixed-effect models fitted for each localizer task within the Multiple Demand network.** The table presents effects of condition, hemisphere, and their interaction for the Language localizer, MD localizer, Articulation, Stroop, and Verbal Fluency tasks.

| **Task** | **Effect** | **Estimate** | **Standard  Error** | ***t* - value** | **Effect size *d*** | ***p*-value** |
| --- | --- | --- | --- | --- | --- | --- |
| **Language localizer** | listening to intact speech > degraded speech | 0,07 | 0,02 | 2,97 | 0,94 | 0,005 |
|  | left > right | -0,02 | 0,03 | -0,54 | -0,20 | 0,596 |
|  | interaction | -0,05 | 0,02 | -2,42 | -0,09 | 0,016 |
| **MD localizer** | hard vWM > easy vWM | -0,47 | 0,05 | -9,36 | -2,96 | 0,000 |
|  | left > right | 0,07 | 0,26 | 0,27 | 0,12 | 0,794 |
|  | interaction | -0,01 | 0,04 | -0,27 | -0,01 | 0,787 |
| **Articulation** | syllable repetition > motor sequence | 0,26 | 0,05 | 5,03 | 1,59 | 0,000 |
|  | left > right | -0,02 | 0,03 | -0,51 | -0,17 | 0,613 |
|  | interaction | -0,05 | 0,03 | -1,71 | -0,06 | 0,088 |
| **Stroop** | naming color words > adjectives | -0,26 | 0,03 | -8,21 | -2,59 | 0,000 |
|  | left > right | -0,19 | 0,08 | -2,30 | -0,98 | 0,031 |
|  | interaction | 0,08 | 0,03 | 2,56 | 0,09 | 0,011 |
| **Verbal Fluency** | fluency > automated speech | -0,04 | 0,06 | -0,72 | -0,23 | 0,476 |
|  | left > right | -0,11 | 0,08 | -1,32 | -0,59 | 0,202 |
|  | interaction | 0,70 | 0,03 | 25,38 | 0,91 | 0,000 |

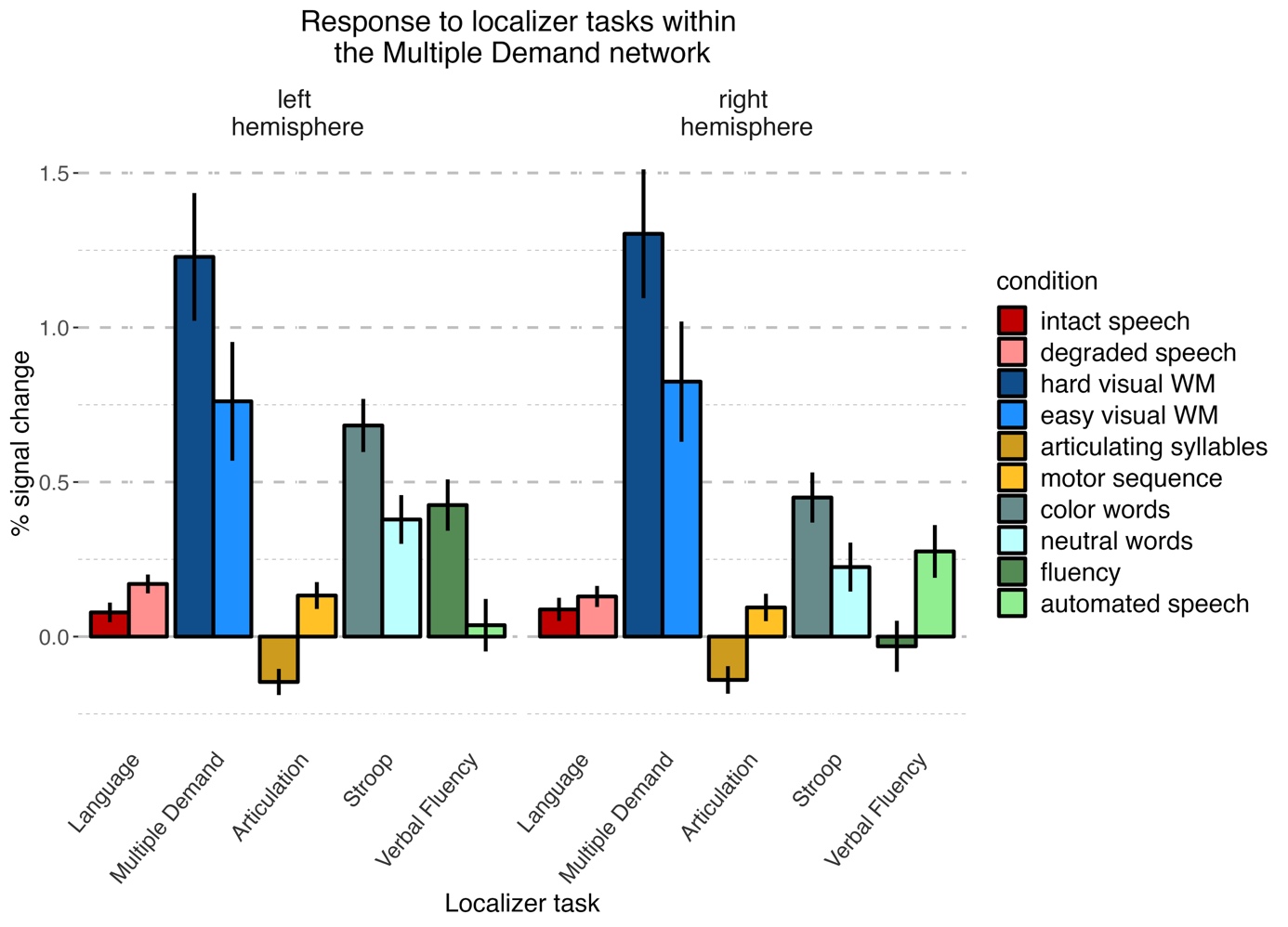

**Figure S7.** **Mean percent signal change corresponding to the localizer task conditions in the Multiple Demand network.** Errorbars represent standard error of the effect of condition in each task and hemisphere.

#### Response to the Language and Multiple Demand localizer tasks in all networks used in this study

For the Language localizer task, we found stronger responses to listening to intact vs. degraded speech in the language and Articulation networks. An opposite effect, i.e., stronger responses to listening to degraded speech vs. intact speech were found in the Multiple Demand and Stroop networks. No differences between the Language localizer conditions were found in the Verbal Fluency network. For the Multiple Demand localizer task, we found stronger responses to hard vs. easy working memory task in the Multiple Demand, Stroop, and Verbal Fluency networks. An opposite effect, i.e., stronger responses to easy vs. hard working memory task was found in the Artuculation network. No differences between the Multiple Demand localizer conditions were found in the language network. The results of these analyses are presented in Tabel S8 and Figure S8.

**Table S8. Results of linear mixed-effect models fitted for the Language and Multiple Demand localizer tasks within five functional networks used in the current study.** The table presents effects of condition for the Language localizer task (listening to intact > degraded speech) and the Multiple Demand localizer task (hard vs easy visual working memory).

| **Functional network** | **Localizer task** | **Estimate** | **Standard  Error** | ***t* - value** | **Effect size *d*** | ***p*-value** |
| --- | --- | --- | --- | --- | --- | --- |
| Language | Language | 0,343 | 0,032 | 10,701 | 3,384 | 0,000 |
|  | Multiple Demand | -0,026 | 0,021 | -1,277 | -0,404 | 0,209 |
| Multiple Demand | Language | -0,067 | 0,023 | -2,974 | -0,941 | 0,005 |
|  | Multiple Demand | 0,473 | 0,050 | 9,363 | 2,961 | 0,000 |
| Articulation | Language | 0,288 | 0,027 | 10,627 | 3,361 | 0,000 |
|  | Multiple Demand | -0,066 | 0,019 | -3,492 | -1,104 | 0,001 |
| Stroop | Language | -0,059 | 0,021 | -2,805 | -0,887 | 0,008 |
|  | Multiple Demand | 0,443 | 0,044 | 10,029 | 3,171 | 0,000 |
| Verbal Fluency | Language | 0,000 | 0,023 | 0,017 | 0,005 | 0,987 |
|  | Multiple Demand | 0,217 | 0,036 | 5,980 | 1,891 | 0,000 |

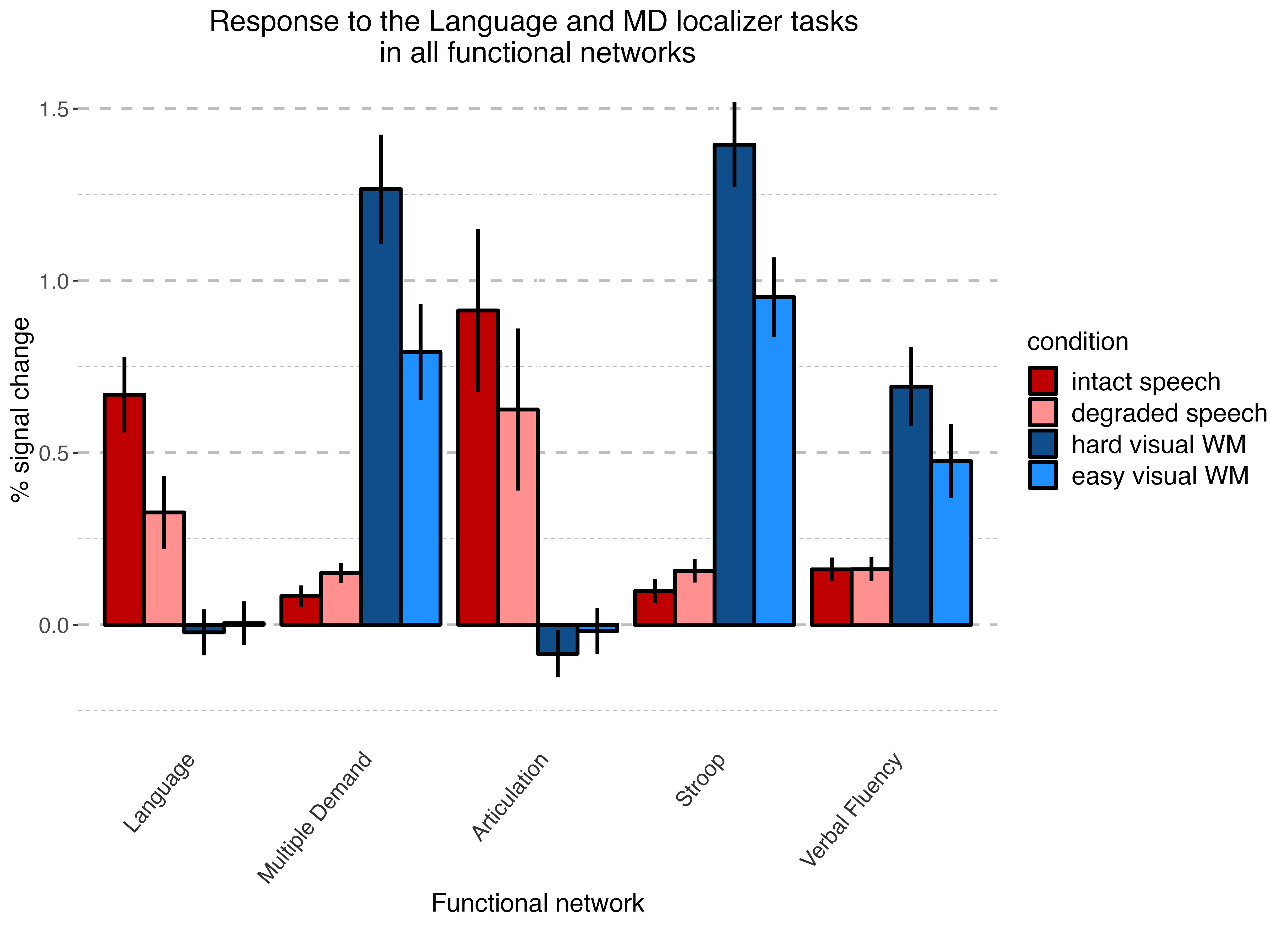

**Figure S8.** **Mean percent signal change corresponding to the Language localizer task and Multiple Demand localizer task conditions in five functional networks used in this study.** Errorbars represent standard error of the effect of condition in each task and hemisphere.

### L2 after-effect in the Anterior Cingulate Cortex

Neurocognitive accounts assume that the Anterior Cingulate Cortex (ACC) is one of the key nodes supporting proactive cognitive control (e.g. Braver et al., 2012). The ACC was not one of the ROIs included in the Multiple Demand network parcels set that we used in this study (Fedorenko et al., 2011), however, to be able to evaluate our results against theories arguing for a crucial role of the ACC in proactive control, we conducted a supplementary analysis in the ACC.

Two ROIs corresponding to the left and right ACC were extracted from the Harvard Oxford Cortical atlas in FSL (Makris et al., 2006; Frazier et al., 2005; Desikan et al., 2006; Goldstein et al., 2007). The ROIs were thresholded at a 25% probability threshold and binarized. Treating the atlas-derived ROIs as the group-level parcels, we subsequently created single-subject ROIs following the group-contrained-subject-specific approach (Fedorenko et al., 2010) which is described in detail in the Data Analysis section of the main manuscript. We used the hard > easy contrast derived from the Multiple Demand localizer to create single-subject ROIs. Finally, we fitted a linear mixed-effect model for each ROI (left and right ACC) using the following formula:

% signal change ~ PrecedingLanguage +

(1 + condition | Subject)

The results revealed no significant effect of condition *(L1 after L1* vs *L1 after L2)* in either the left ACC ROI (*t(40)* = 0.171, *p* = 0.865) or the right ACC ROI (*t(40)* = 0.694, *p* = 0.492). The results are presented in **Figure S9.**

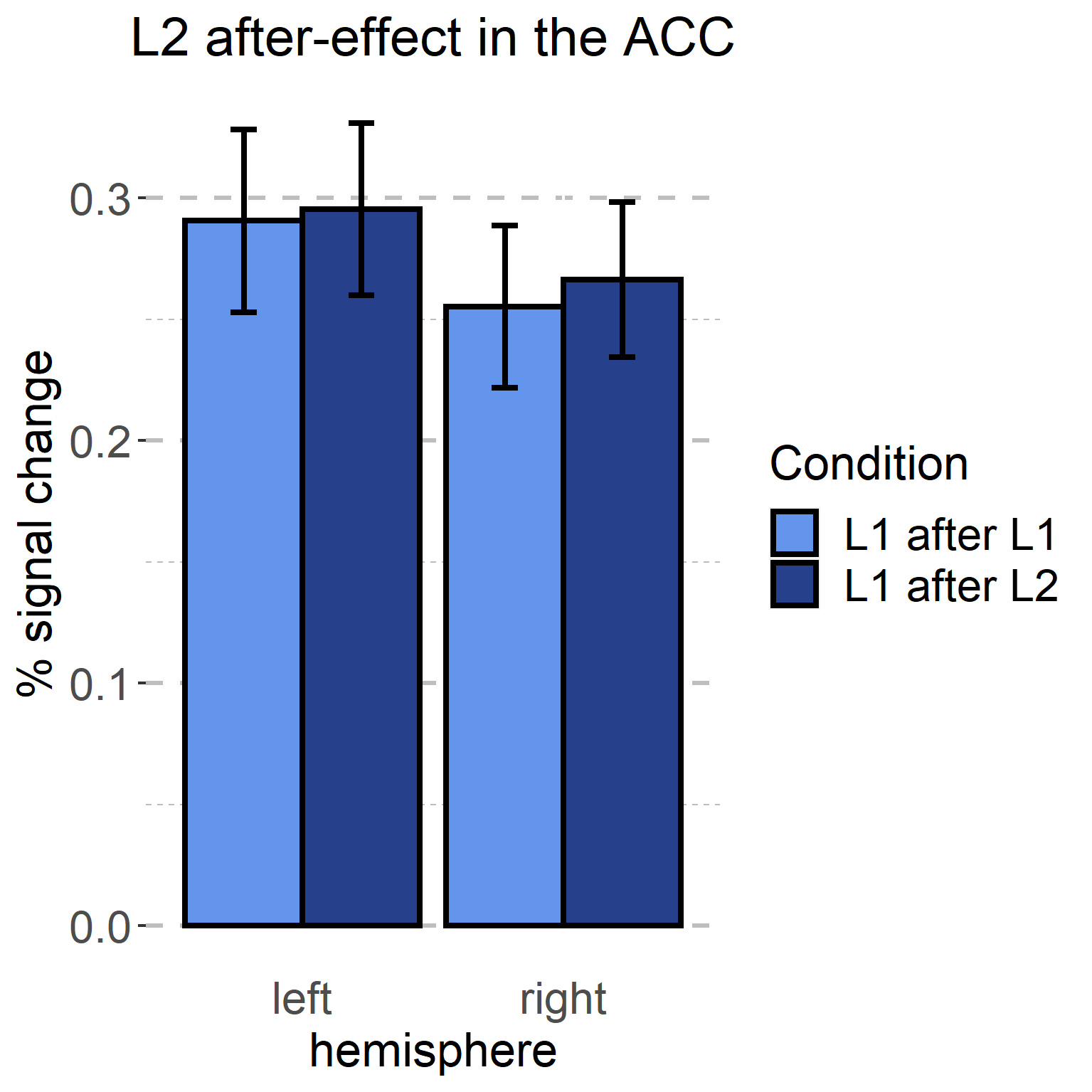

**Figure S9.** Mean % signal change for the *L2 after-effect* in the left and right ACC ROIs. Whiskers correspond to the standard error of the mean.

### Behavioural data analysis and results

#### Data analysis

Behavioral naming data recorded in the scanner were transcribed for each participant and analyzed for naming latencies and accuracy. One participant’s data were excluded because of very poor recording quality that did not allow us to determine the naming latencies in one of the sessions^[1]^. For the picture naming task, all answers corresponding to the dominant response and its synonyms were considered correct. Trials with no responses, inaudible responses, and names unrelated to a given picture were classified as errors and excluded from further analysis of naming latencies. The naming latencies for each answer were determined manually. Subsequently, we fitted a linear mixed-effect model to the naming latency data and a generalized mixed-effect linear model to the accuracy data (using the lmer() and glmer() functions from the lmerTest (Kuznetsova et al., 2017) and lme4 package (Bates et al., 2015), respectively). As it has been shown (Casado et al., 2022), the behavioral *L2 after-effect* largely depends on a language balance of a given speaker, i.e., it is larger for bilinguals for whom the difference between L2 and L1 activation (i.e., those that name pictures faster in their L1 than in L2). To take this into account, we included a predictor of language balance (i.e., difference between L1 and L2 naming latencies measured in the first task blocks) into the behavioral analysis of the L2 after-effect. We used the following formula:

Naming latencies / Accuracy ~ PrecedingLanguage + LanguageBalance + PrecedingLanguage:LanguageBalance +

(1 + PrecedingLanguage | Subject) +

(1 + PrecedingLanguage + LanguageBalance + PrecedingLanguage:LanguageBalance | Item)

*PrecedingLanguage* corresponded to a predictor with two levels: *L1 after L1* and *L1 after L2*. *LanguageBalance* corresponded to the difference between mean naming latencies in L1 and L2 measured in the first block of picture naming task in each session (for reference, see Figure 1). The categorical predictor was deviation-coded (L1 after L1 = -0.5; L1 after L2 = 0.5); the continuous predictor of language balance was demeaned and the naming latencies were log-transformed to reduce the skewness of the data (skewness before and after transformation was equal to 0.80 and 0.26, respectively). For each analysis, we first fitted a maximal model and then identified the maximum random-effects structure justified by the data, following the recommendations of Bates et al. (2018).

#### Results

#### Behavioral results

The analysis of naming latencies revealed no main effect of preceding language (β = 0.005, *t* = 0.54, *p* = .590) or balance (β = -0.015, *t* = -1.31, *p* =.198) but it revealed a significant interaction of preceding language and balance (β = 0.045, *t* = 4.95, *p* < 0.001), showing that the L2 after-effect was larger for more unbalanced speakers (see Figure S10). Mean naming latencies predicted by the model corresponded to 1070ms (CI [1043ms, 1099]) for *L1 after* L1 and 1076 (CI [1048, 1105ms]) for *L1 after* L2. Similarly, accuracy analysis did not reveal an effect of preceding language (β = -1.015, *z* = -1.437, *p* = .151) or balance (β = 0.272, *z* = 1.660, *p* = .97). Unlike the naming latencies, accuracy results did not reveal a significant interaction between the preceding language and balance (β = -0.551, *z* = -1.69, *p* = 0.091). Accuracies predicted by the model correspond to 99.6% (CI [98.65%, 99.88%]) for the *L1 after L1* condition, and 98.89% (CI [98.12%, 99.34%]) for the *L1 after L2* condition*.* In line with previous reports (Casado et al., 2022), we found that the *L2 after-effect* was modulated by the balance between L1 and L2: the difference between naming latencies in L1 after L1 and L1 after L2 was larger for participants who were much slower in naming pictures in L2 than L1 (i.e., they were less balanced). The current results also replicate our previous finding that after L2, L1 slow-down is similar regardless of the balance; however, in more balanced bilinguals the baseline L1 (i.e., the L1 after L1 condition) is overall slower than in unbalanced, L1-dominant bilinguals (see Figure 5, Casado et al., 2022). It is also important to note that even though participants were instructed to name pictures as quickly as possible, overall naming speed (1072ms for *L1 after L1* and 1075ms for *L1 after L2*) was slower than what is usually observed for L1 naming for this particular picture set (897ms; Wolna et al., 2022). This may be because we collected the behavioral data during scanning, in a generally noisy and challenging environment. Still, despite the challenges related to overt naming data collection in the scanner, we replicated the relationship between L2 after effect and language balance. One important implication for the neuroimaging data analysis is that the *L2 after-effect* may not affect balanced participants, as they do not experience the behavioral slow-down. To account for this possibility, additionally to the localizer-based analyses on the whole group, we have run an additional analysis on a subset of data from 31 clearly unbalanced subjects (defined as having the difference between mean naming speed in L2 vs. L1 larger than 50 ms). However, the analyses on the unbalanced sub-group yielded qualitatively the same results as those based on the whole group. We found significant difference beteen brain response *to L1 after L1* and *L1 after L2* in the Multiple Demand network (β = 0.680, *t* = 2.26, *p* = .031) but no significant differences were found in the Language network (β = 0.042, *t* = 1.40, *p* = .180), Articulation network (β = 0.023, *t* = 0.48, *p* = .363), Stroop network (β = 0.024, *t* = 0.625, *p* = .537), or Verbal Fluency network (β = -0.030, *t* = -0.77, *p* = .446).

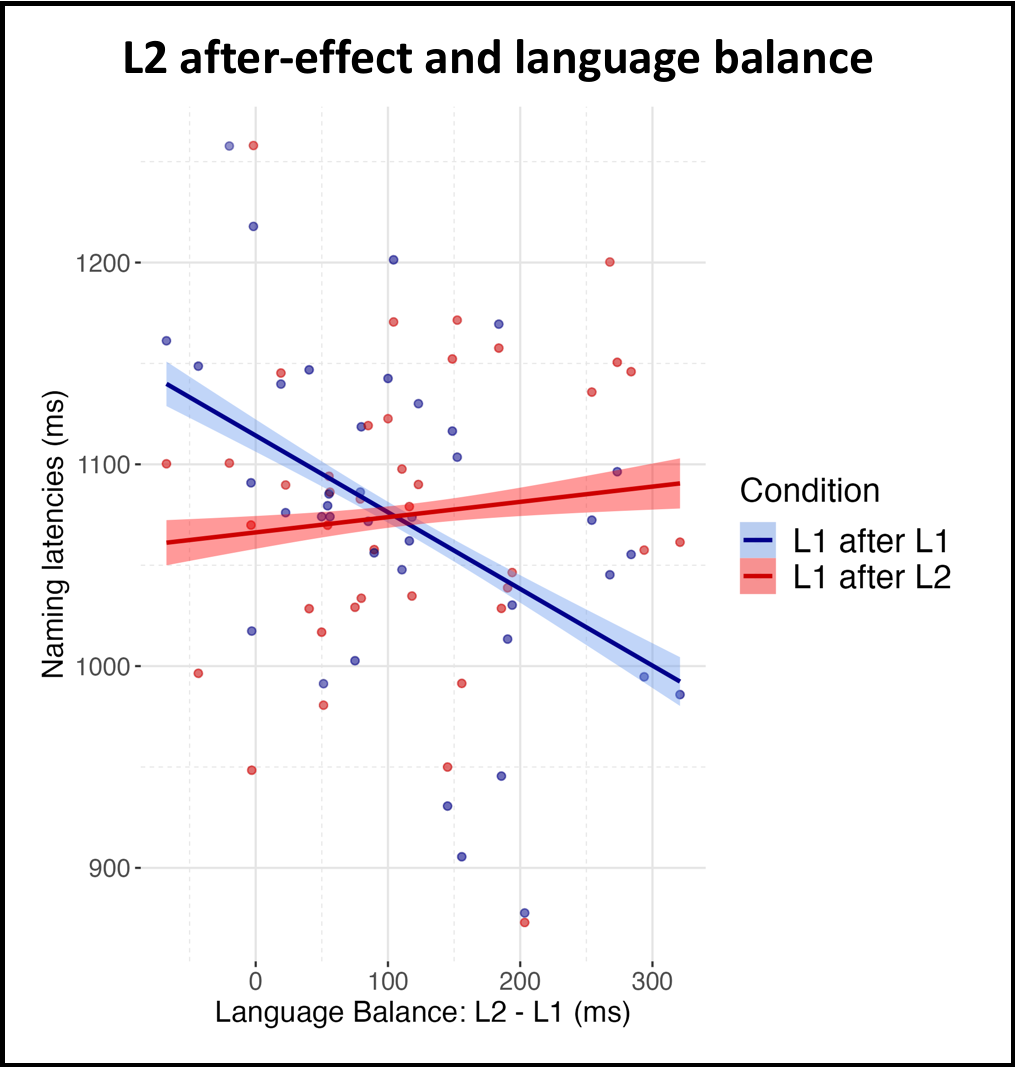

**Figure S10.** Results of the analysis of naming latencies in *L1 after L1* and *L1 after L2* depending on the participant’s balance. Individual data points correspond to mean predicted naming latencies for each condition and each subject

### Lists of stimuli

#### Verbal fluency task

The verbal fluency task used as a functional localizer was performed only in the participants’ L1. Below we provide a full list of stimuli for both the phonemic and semantic conditions (along with a translation of the latter).

| **task** | **cue** | **translation** |
| --- | --- | --- |
| category | [masked for review] | geometric figures |
| category | [masked for review] | names of flowers |
| category | [masked for review] | European countries |
| category | [masked for review] | clothes |
| category | [masked for review] | vegetables |
| category | [masked for review] | body parts |
| category | [masked for review] | tree species |
| category | [masked for review] | colors |
| category | [masked for review] | furniture |
| category | [masked for review] | means of transportation |
| category | [masked for review] | sweets |
| category | [masked for review] | car brands |
| letter | G |  |
| letter | D |  |
| letter | H |  |
| letter | K |  |
| letter | L |  |
| letter | W |  |
| letter | A |  |
| letter | C |  |
| letter | P |  |
| letter | R |  |
| letter | T |  |
| letter | Z |  |

1. As this problem was limited to the behavioral data, this participant was still included in the neuroimaging analyses.
